# Conservation of gene architecture and domains amidst sequence divergence in the *hsrω* lncRNA gene across the *Drosophila* genus: An *in silico* analysis

**DOI:** 10.1101/695486

**Authors:** Ranjan Kumar Sahu, Eshita Mutt, Subhash Chandra Lakhotia

**Affiliations:** Cytogenetics Laboratory, Department of Zoology, Banaras Hindu University, Varanasi, 221005 INDIA; Laboratory of Biomolecular Research, Paul Scherrer Institute, 5232 Villigen, Switzerland

**Author notes:** Author emails: Ranjan Kumar Sahu, Eshita Mutt, S. C. Lakhotia.

**Keywords:** *lncRNA*, *ORFω*, *omega* intron, *mir-4951*, *splice junction*, *tandem repeats*

## Abstract

The developmentally active and cell-stress responsive *hsr*ω locus in *Drosophila melanogaster* carries two exons, one *omega* intron, one short translatable open reading frame ORFω, long stretch of unique tandem repeats and an overlapping *mir-4951* near its 3’ end. It produces multiple lncRNAs using two transcription start and four termination sites. Earlier studies revealed functional conservation in several *Drosophila* species but with little sequence conservation, in three experimentally examined species, of ORFω, tandem repeat and other regions but ultra-conservation of 16nt at 5’ and 60nt at 3’ splice-junctions of the *omega* intron. Present bioinformatic study, using the splice-junction landmarks in *Drosophila melanogaster hsr*ω, identified orthologues in publicly available 34 *Drosophila* species genomes. Each orthologue carries the short ORFω, ultra-conserved splice junctions of *omega* intron, repeat region, conserved 3’-end located *mir-4951*, and syntenic neighbours. Multiple copies of conserved nonamer motifs are seen in the tandem repeat region, despite a high variability in repeat sequences. Intriguingly, only the intron sequences in different species show evolutionary relationships matching the general phylogenetic history in the genus. Search in other known insect genomes did not reveal sequence homology although a locus with similar functional properties is suggested in *Chironomus* and *Ceratitis* species. Amidst the high sequence divergence, the conserved organization of exons, ORFω and *omega* intron in this gene’s proximal part and tandem repeats in distal part across the *Drosophila* genus is remarkable and possibly reflects functional importance of higher order structure of *hsr*ω lncRNAs and the small Omega peptide.

## Introduction

Major parts of genomes in diverse eukaryotes are comprised of the non-coding (nc) DNA which exhibit very widely varying evolutionary conservation ranging from ‘ultra-conservation’ across phylogenetic groups to remarkable variability even between related species. A large part of ncDNA is transcribed as different types of short and long non-coding RNAs (sncRNA and lncRNA). Although in the past the nc genomic sequences were often ignored as ‘selfish’ or ‘junk’ DNA, recent years have witnessed a remarkable interest in the ncRNAs because of their well-documented pervasive involvement in cellular regulatory networks in diverse ways and at different levels, ranging from regulation of local chromatin organization, transcription, post-transcriptional processing, transport and stability of RNAs to modulation of activities of different regulatory proteins in the cell (Anastasiadou *et al*., 2018; Fatima *et al*., 2015; Ferreira & Esteller, 2018; Hirose & Nakagawa, 2016; Kopp & Mendell, 2018; Lakhotia, 1996; Lakhotia, 2012; Lakhotia, 2015; Lakhotia, 2016; Lakhotia, 2017a; Lakhotia, 2017b; Lakhotia, 2018; Lakhotia *et al*., 2020; Matsumoto & Nakayama, 2018; Noh *et al*., 2018; Quinn *et al*., 2016; Ransohoff *et al*., 2018; Stoiber *et al*., 2016).

The *93D* or *hsr*ω in *Drosophila melanogaster* (*D. mel*), one of the first identified lncRNA genes, is known to be essential for normal development as well as for surviving cell stress (Lakhotia, 1987; Lakhotia, 2011; Lakhotia, 2012; Lakhotia, 2017a; Lakhotia & Mukherjee, 1982; Mohler & Pardue, 1984). Interest in the stress inducible and developmentally active *93D* locus of *D. mel*, later renamed as *hsr*ω gene (Bendena *et al*., 1989), was generated following a serendipitous observation that the 93D puff was singularly activated after a brief in vitro exposure of larval salivary glands to 1mM benzamide, which otherwise inhibited the on-going transcription at other chromosomal sites (Lakhotia & Mukherjee, 1970). It was found to be one of the most active heat shock induced genes with unusual turnover of its transcripts (Lakhotia & Mukherjee, 1980; Lengyel *et al*., 1980; Mukherjee & Lakhotia, 1979; Peters *et al*., 1980), which were apparently not translated (Lakhotia & Mukherjee, 1982).

Cloning and sequencing of this locus from three distantly related species, viz., *D. mel, D. hydei* (*D. hyd*) and *D. pseudoobscura* (*D. pse*) (Garbe *et al*., 1986; Garbe *et al*., 1989; Garbe & Pardue, 1986; Peters *et al*., 1984) confirmed its non-coding nature and revealed a comparable organization of the gene although with high sequence variability. Subsequent studies established a critical importance of this gene in normal development as well as in cell stress response (Lakhotia, 1987; Lakhotia, 2011). This gene is transcribed in nearly every cell type in the life of fly (Bendena *et al*., 1991; Lakhotia *et al*., 2001; Mutsuddi & Lakhotia, 1995). Of its multiple transcripts, the repeat containing nuclear transcripts are essential for organization of the nucleoplasmic omega speckles, which coordinate the dynamic availability of diverse hnRNPs and some other RNA binding proteins in cells under normal and stress conditions (Lakhotia *et al*., 2012; Lakhotia *et al*., 1999; Lo Piccolo, 2018; Mallik & Lakhotia, 2009; Mallik & Lakhotia, 2011; Onorati *et al*., 2011; Prasanth *et al*., 2000; Singh & Lakhotia, 2015).

The *hsr*ω gene in *D. mel* is located at the 93D4-5 cytogenetic region on right arm of chromosome 3 (genomic coordinates 3R:21,296,127…21,317,836, http://flybase.org/reports/FBgn0001234). Following the advent of genome sequencing, the annotated genomes of 5 *Drosophila* species (including *D. mel*) are available at the FlyBase (http://flybase.org/cgi-bin/gbrowse2/) while unannotated genome sequences of many more species are available at the NCBI database (https://www.ncbi.nlm.nih.gov/). Early studies (Garbe *et al*., 1986; Lakhotia, 1987; Peters *et al*., 1982; Peters *et al*., 1984) suggested that this gene has common architectural features with ∼10 to 15kb long transcription unit, whose proximal part (∼2.6kb) comprises of unique sequence while the distal region (>5kb) contains tandem repeats of a short sequence, unique to this locus. Intriguingly, the unique part did not show any significant sequence homology between *D. mel, D. hyd* and *D. pse* except for remarkably conserved stretches of 16 and 60 nucleotide at the 5′ and 3′ splice junctions, respectively, of the single intron, which we now term as the *omega* intron. The other identified conserved region was multiple copies of nonamer motif (ATAGGTAGG) in the repeat part of the large nuclear transcript in *D. mel* and *D. hyd* (Garbe *et al*., 1986). These early studies indicated this gene to produce two independent primary nucleus limited transcripts, with the >10kb longer (*hsr*ω*-n*) transcript corresponding to the entire transcription unit and the independently transcribed shorter 1.9kb (*hsr*ω*-pre-c*) transcript being derived from the proximal unique region till the first poly-A signal located about 700 bp upstream of the beginning of the stretch of tandem repeats. Splicing of the single intron (∼700 bases long) in the 1.9kb nuclear transcript produced the 1.2kb (*hsr*ω*-c*) cytoplasmic transcript, which carried a potentially translatable open reading frame (ORFω) that could encode 23–27 amino acids in the three examined species (Fini *et al*., 1989; Garbe *et al*., 1986). Intriguingly, the amino acid sequence potentially encoded by the ORFω in the three species did not appear to show high conservation (Garbe *et al*., 1986). Absence of strong conservation of its base sequence in the three examined species of *Drosophila* (Garbe *et al*., 1986; Garbe *et al*., 1989; Garbe & Pardue, 1986; Peters *et al*., 1984; Ryseck *et al*., 1987) appear rather incongruous with its known (Lakhotia, 1987; Lakhotia, 2011; Lakhotia, 2016) critical functions in cell regulation.

Currently the NCBI and FlyBase databases annotate the *hsr*ω gene of *D. mel* to produce 7 transcripts (Fig. 1). Among these, the *hsr*ω*-RA* (*hsr*ω*-c*), *hsr*ω-*RB* (*hsr*ω*-n1*), *hsr*ω-*RC* (*hsr*ω*-pre-c*) and *hsr*ω-*RG* (*hsr*ω*-n2*) transcripts have been known earlier (Lakhotia, 2011), while the *hsr*ω-*RD, hsr*ω-*RH*, and *hsr*ω-*RF* transcripts are new additions on the basis of RNA-sequence data. These seven transcript variants originate because of different combinations of 2 transcription start sites (TSS), 4 transcription termination sites (TTS) and variable splicing of the single intron (see Fig. 1). The TSS for *hsr*ω-*RD* is named as TSS1 while that for all the other four primary transcripts, viz., *hsr*ω-*RB* (*hsr*ω*-n1*), *hsr*ω-*RC* (*hsr*ω*-pre-c*), *hsr*ω-*RF* and *hsr*ω-*RH*, located 494 bp downstream of TSS1, is named TSS2 (http://flybase.org/reports/FBgn0001234). The FlyBase report for this gene states that orthologs of this gene are not known in other *Drosophila* species or in any other organism.

**Fig. 1.**
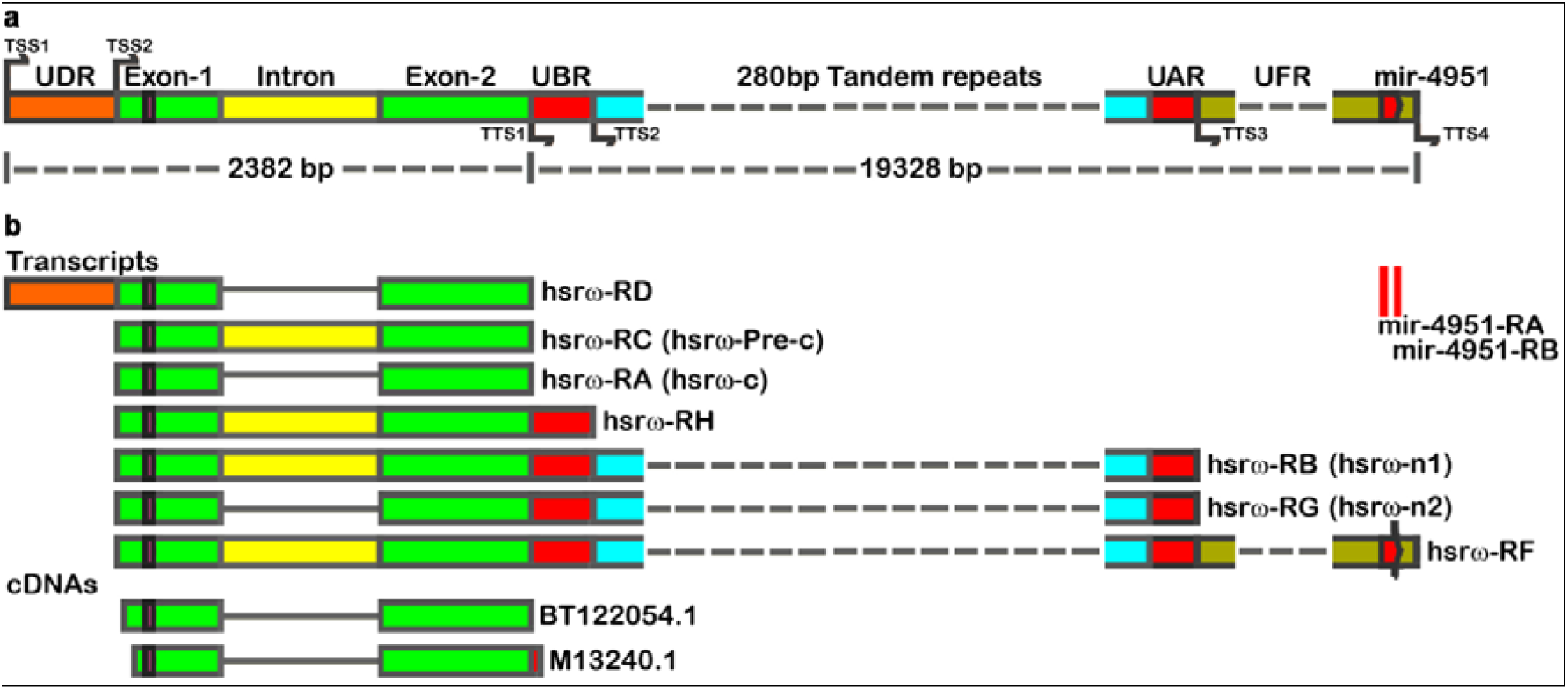
Genomic coordinates and predicted transcripts of the *hsr*ω gene in *D. mel* (http://flybase.org/reports/FBgn0001234). **a**. Organization of the *hsr*ω gene as available at FlyBase: two upward arrows in indicate the two transcription start sites (TSS1 and TSS2) while the four downward arrows indicate the four annotated transcription termination sites (TTS1-TTS4). **b**. Transcripts produced by the *hsr*ω gene: unspliced intron in transcripts is shown as yellow box while the spliced out intron is indicated by a black line joining the exon 1 and exon 2 regions (green boxes). UDR = region unique to RD transcript, UBR = unique region before tandem repeats between TTS1 and TTS2, UAR = unique region after tandem repeats till the TTS3; UFR = unique region specific to the RF transcript, *mir-4951* = miRNA gene overlapping with the *hsr*ω gene at its 3’ end, broken lines indicate the 280 bp tandem repeats (redrawn after http://flybase.org/reports/FBgn0001234).

Comparative genomic analysis of multiple genomes has remarkably improved the resolution for analysis of phylogenetic relationships among different species. Since information about genome sequence for a large number of *Drosophila* species is now available in public databases and since the above noted earlier studies indicated a variable conservation of the *hsr*ω lncRNA gene, we undertook the present *in silico* analysis of base sequence and architecture of the *hsr*ω gene in 34 species of *Drosophila*, viz., *D. albomicans* (*D. alb*), *D. americana* (*D. ame*), *D. ananassae* (*D. ana*), *D. arizoanae* (*D. ari*), *D. biarmipes* (*D. bia*), *D. bipectinata* (*D. bip*), *D. busckii* (*D. bus*), *D. elegans* (*D. ele*), *D. erecta* (*D. ere*), *D. eugracilis* (*D. eug*), *D. ficusphila* (*D. fic*), *D. grimshawi* (*D. gri*), *D. hydei* (*D. hyd*), *D. kikkawai* (*D. kik*), *D. melanogaster* (*D. mel*), *D. miranda* (*D. mir*), *D. mojavensis* (*D. moj*), *D. montana* (*D. mon*), *D. nasuta* (*D. nas*), *D. navojoa* (*D. nav*), *D. novamexicana* (*D. nov*), *D. obscura* (*D. obs*), *D. persimilis* (*D. per*), *D. pseudoobscura* (*D. pse*), *D. rhopaloa* (*D. rho*), *D. sechellia* (*D. sec*), *D. serrata* (*D. ser*), *D. simulans* (*D. sim*), *D. subobscura* (*D. sub*), *D. suzuki* (*D. suz*), *D. takahashii* (*D. tak*), *D. virilis* (*D. vir*), *D. willistoni* (*D. wil*), and *D. yakuba* (*D. yak*). The 34 species examined in our study belong to 8 sub-genus/species groups, viz., *melanogaster* (16 species), *repleta* (4 species), *Hawaiian* (1 species), *immigrans* (2 species), *obscura* (5 species), *virilis* (4 species), *willistoni* (1 species) *and Dorsilopha* (1 species) (O’Grady & DeSalle, 2018). For convenience, the 34 species are referred to in the following using only the first three letters (as in parentheses above) of their species names.

The *hsr*ω lncRNA gene has been studied and annotated extensively in *D. melanogaster*, while some information about its structure and expression patterns in a few other *Drosophila* species is also available. However, its genomic organization has not yet been annotated in any other species. Present study provides basic genomic features of the *hsr*ω lncRNA gene in 33 other *Drosophila* species. We show that some features of the architecture of this long gene are remarkably conserved in all the 34 *Drosophila* species examined although its base sequence shows a peppered conservation with only certain small regions being significantly similar between the 34 species. Contrary to the generally known faster divergence of intronic than exonic sequences in related species, the evolutionary divergence of the omega intron sequence parallels the known phylogenetic relationships of different *Drosophila* species, while diversification of the exonic and other regions does not correlate with the phylogenetic relationships of different species. The only other region whose sequence divergence between species partially matches their phylogenetic relationship is the ORFω. This first-time annotation of the *hsr*ω lncRNA gene in large number of *Drosophila* species would be very useful for future functional studies on this developmentally important and cell-stress responsive gene.

## Materials and Methods

### Retrieval of genomic sequences carrying *hsr*ω orthologue in different species

Sequences flanking the 5’ (16 bp) and 3’ (60 bp) splice junctions were shown in earlier studies (Garbe *et al*., 1986) to be nearly completely identical between *D. mel, D. hyd* and *D. pse* species. Therefore, we used these as query sequences to search for homologous stretches in the other 33 *Drosophila* species’ genome sequences available, as on 17 Nov 2018, at the NCBI and FlyBase genome databases (12 species being common in NCBI and FB2017_05 October (Dmel Release 6.18) using the online BLAST NCBI (http://www.ncbi.nlm.nih.gov/BLAST/) and BLAST FlyBase (http://flybase.org/blast/) programs. Only the genome sequences that showed presence of both the 5’ (16 bp) and 3’ (60 bp) splice junction sequences in neighbourhood were selected for further analysis.

The *hsr*ω gene in *D. mel* carries a miRNA gene, *mir-4951*, located ∼971 bp upstream of the annotated TTS4 (Fig. 1) at the end of its largest transcript *RF* (http://flybase.org/reports/FBgn0001234). Our initial survey revealed that this miRNA sequence is present downstream of the *hsr*ω orthologue in all the other 11 species whose genomes are available at the FlyBase. Therefore, we queried all the other 33 species’ genomes with the *mir-4951* sequence of *D. mel* to retrieve its flanking sequences.

The above two sets of retrieved sequences were aligned with Refseq Genome Database (Refseq_Genomes) and Whole-genome shotgun contigs (WGS) databases. In most cases, we could retrieve at least 40kb of contiguous genomic sequences (see Results) that covered 2kb upstream and 38kb downstream of the identified 5’ splice junction, and used the same for further analyses.

### Sequence annotation and mapping

An offline tool SnapGene® (GSL Biotech, <u>snapgene.com</u>) was used for sequence annotation and mapping of different regions of the *hsr*ω genes of the 34 *Drosophila* species.

### Splice site prediction

The region starting with GT (12th and 13th nucleotide (nt) in the 16 bp 5’ junction and ending with AG (39th and 40th nt) in the 60 bp 3’ junction (Garbe *et al*., 1986) in the genomic sequences retrieved for each species was annotated as intron. This was further confirmed by BDGP Splice Site Prediction by Neural Network tool (http://www.fruitfly.org/seq_tools/splice.html) and Net Gene 2 predictor program (http://www.cbs.dtu.dk/services/NetGene2/) using the retrieved sequences from 34 *Drosophila* species genomes as input. Sequences were searched against *Drosophila* as well as *Human* splice site database for the forward strand with a minimum score of 0.4.

### Promoter and TSS prediction

Since the *hsr*ω gene of *D. mel* has two transcription start sites (TSS) in the proximal region of the gene (Fig. 1), we used the BDGP Neural Network Promoter Prediction program (http://www.fruitfly.org/seq_tools/promoter.html) to search for potential promoter and TSS in 2kb region upstream of the identified 5’ splice junction in the retrieved genomic sequences. A minimum score of 0.8 was set as default for search against eukaryotic promoter database with search restricted to the forward strand of the putative *hsr*ω gene in each species.

### PolyA and 3’ end cleavage site prediction

The polyadenylation and transcription termination sites (TTS) in the retrieved sequences were searched using the online tool PolyAh, which recognizes 3’-end cleavage and polyadenylation region (Salamov & Solovyev, 1997) (http://www.softberry.com/berry.phtml?topic=polyah&group=help&subgroup=promoter).

### Search for ORFs, including the ORFω orthologue

ORF finder (https://www.ncbi.nlm.nih.gov/orffinder/) and SnapGene® tools were used to search for potential orthologue of the *D. mel* ORFω in the retrieved putative *hsr*ω genomic sequences of other species using default parameters that the potential ORF must start with ATG, must be coding for at least 20 amino acids, should be in forward orientation and lie in between TSS2 and 5’ splice junction.

### Protein secondary structure prediction

In order to study the structural similarities of the putative peptide produced by the ORFω, the predicted amino acid sequences were analysed for their secondary structures using PSIPRED (Jones, 1999) (http://bioinf.cs.ucl.ac.uk/psipred/) tool.

### Estimation of repetitive sequences

The length and frequency of tandem repeats in the retrieved genomic sequences were analysed using the Tandem Repeat Finder tool (Benson, 1999) (http://tandem.bu.edu/trf/trf.html) with a minimum period size set to 7 bp and maximum to 500 bp.

### Evolutionary conservation analysis

Nucleotide and amino acid sequences from different regions of *hsr*ω gene in different species were aligned and percent identity matrix (PIM) was calculated with help of the multiple sequence alignment (MSA) tool T-Coffee (https://www.ebi.ac.uk/Tools/msa/tcoffee/), which provided conservation colour codes as BAD AVG GOOD; these were used for all alignment tables and figures. Evolutionary relationships were inferred using the Maximum Likelihood method with Mega X software (Kumar *et al*., 2018).

## Results

### Early cytogenetic identification of hsrω orthologs in different species of Drosophila

The unique induction of one of the major heat shock puffs in *D. mel* with benzamide (Lakhotia & Mukherjee, 1970) (Mukherjee & Lakhotia, 1982), presence of large RNP particles at this puff (Dangli *et al*., 1983) and accumulation of several hnRNPs at this site during heat shock (Saumweber *et al*., 1980) have been utilized in early cytogenetic studies to identify an orthologue of *93D* or *hsr*ω gene in several species groups of *Drosophila*. In *D. hyd* and other species, the same locus (2-48C) was also known to be selectively induced by brief in vitro exposure of larval salivary glands to vit-B6 or pyridoxal phosphate (Leenders *et al*., 1973), although vit-B6 failed to induce any puff in *D. mel* (Lakhotia & Singh, 1982). Table 1 summarizes the earlier cytogenetic evidence for existence of *hsr*ω orthologue in many *Drosophila* species. These studies showed that a gene sharing the unique inducible properties of *D. mel hsr*ω was associated, in all the examined species, with ancestral chromosome element E (Muller, 1940). These cytogenetic studies clearly suggested that a functional orthologue of *hsr*ω gene is present in diverse *Drosophila* species although results of the few early DNA:DNA or RNA:RNA in situ hybridization studies (Table 1) and direct sequencing of the cloned regions of this gene in a few species (Drosopoulou *et al*., 1996; Garbe *et al*., 1986; Garbe *et al*., 1989; Peters *et al*., 1984) revealed a rather limited conservation of DNA sequence at this locus.

**Table 1.**
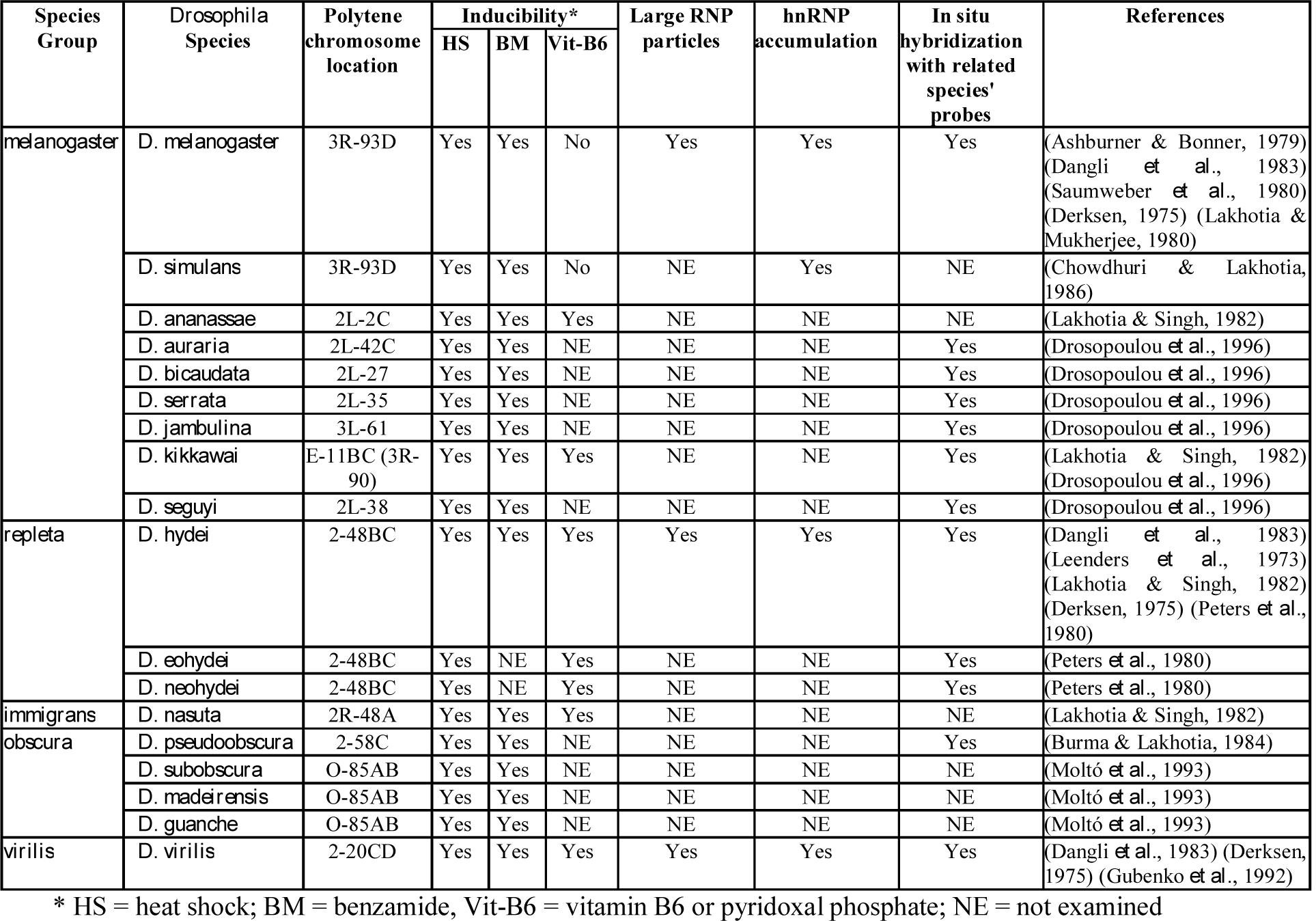
Cytogenetic identification of *hsr*ω-like loci in different species of *Drosophila*.

### Genomes of all Drosophila species carry hsrω orthologue and show a conserved order of flanking genes

Our search of genomes of the 34 *Drosophila* species for the 16 bp 5’ and 60 bp 3’ splice junction associated sequences, and for the *mir-4951* sequence revealed that the two splice junction associated sequences were found in neighbourhood only once in each of the 34 species genomes. In 27 species, the retrieved sequences also carried the *mir-4951* orthologue downstream of the 3’ splice junction sequence. In the case of *D. sec*, we used its recently assembled complete genome sequence (Chakraborty *et al*., 2020) to blast and align the two separate scaffolds retrieved from the databases. This revealed that the two identified scaffolds flanked a region (∼19kb) reported in the genome sequence of *D. sec* (Chakraborty *et al*., 2020). The resulting contig showed presence of the *mir-4951* orthologue downstream of the 3’ splice junction sequence. In seven other species (*D. ame, D. eug, D. moj, D. nas, D. obs, D. rho*, and *D. suz*), whose genomes are not yet fully assembled, the retrieved scaffolds carried either the two splice junctions or the *mir-4951* sequence. Significantly, the region downstream of the 3’ splice junction or upstream of the *mir-4951* in the identified scaffolds in all these seven cases was small (Fig. 2). The scaffold that carried the *mir-4951* orthologue was blasted with that species’ scaffold carrying the 5’ and 3’ splice junctions to find any overlap. In the case of *D. ame* and *D. suz*, the splice junctions, and the *mir-4951* sequence carrying scaffolds showed overlapping regions (52 bp in *D. ame* and 146 bp in *D. suz*), using which we assembled contigs for the two species. In the other five species (*D. eug, D. moj, D. nas, D. obs* and *D. rho*), the two scaffolds did not show any overlap with each other (Fig. 2), or common overlap with any other scaffold. Consequently, the complete *hsr*ω region could not be assembled for these five species. A contiguous stretch of genomic sequence, at least 2kb upstream and 38kb downstream of the identified 5’ splice junction, was used for further analyses in the 29 species while for the other five species, the two separate scaffolds, carrying the splice junctions and *mir-4951* sequence, respectively, were used.

**Fig. 2.**
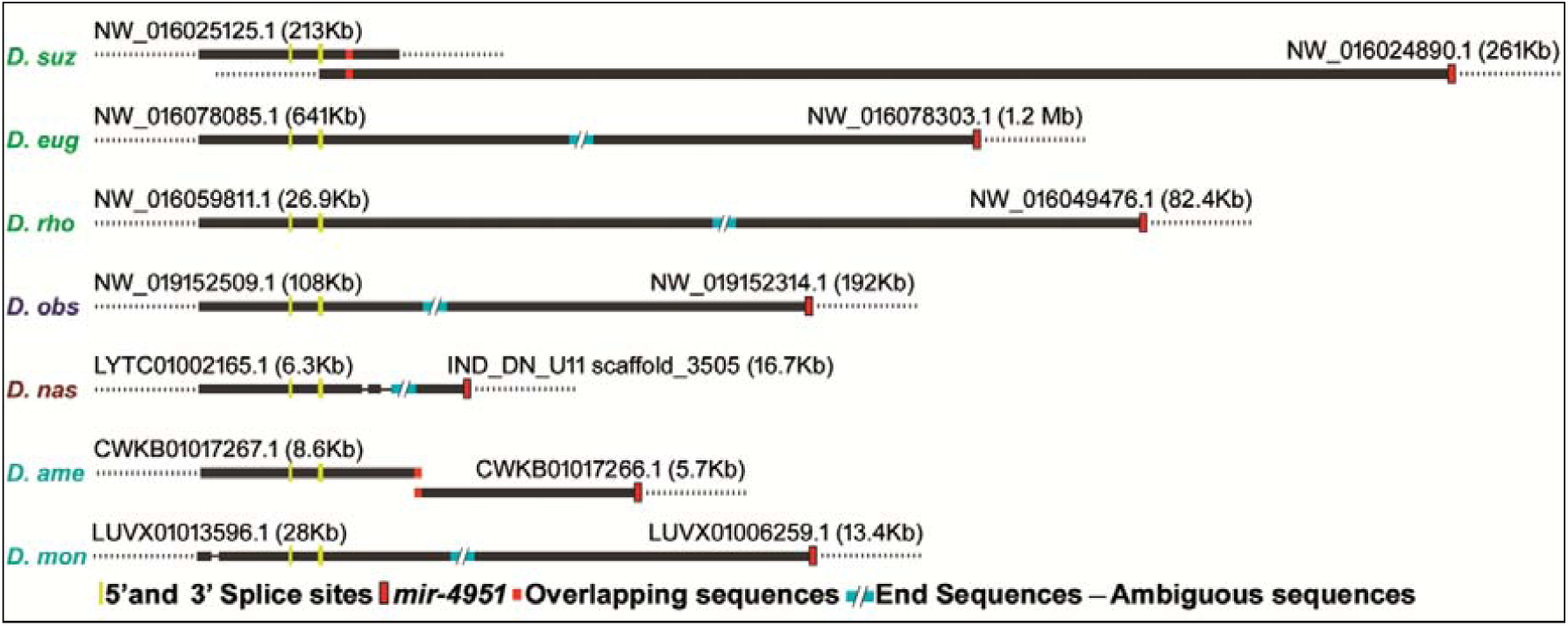
Strategy for assembly of *hsr*ω carrying contigs in species with the splice junctions and *mir-4951* retrieved in separate scaffolds. Schematics of the scaffolds carrying splice junctions or the *mir-4951* sequences in different species are shown as solid thick (putative *hsr*ω regions) and dotted lines (flanking regions). The yellow and red vertical bars represent the splice junctions and the *mir-4951* sequences, respectively; orange vertical bars indicate regions of sequence overlap between the two scaffolds in *D. ame* and *D. suz*, while slanted turquoise bars represent non-overlapping but presumably facing ends of the two scaffolds in six species.

We aligned (Fig. 3) the retrieved sequences/scaffolds with the partially annotated genome data for the 4 other species (*D. ana, D. sim, D. pse*, and *D. vir*) as available at the current version of FlyBase (FB2020_01). The retrieved sequences could be associated with annotated genes in all of these species, which we rename as the *hsr*ω gene (Fig. 3). To confirm identity of the above retrieved genomic sequences with *D. mel hsr*ω, we examined if genes known to flank the *hsr*ω locus in *D. mel* also flanked the genomic sequences retrieved for the other 33 species. The *hsr*ω locus in *D. mel* is flanked on upstream by *SIFaR* and *ETHR* genes and on downstream by *CG16791, mod, tin* and *bap* genes (http://flybase.org/reports/FBgn0001234). Sequences of these *D. mel* genes were blasted with the genomes of the other 33 species and their association with the retrieved putative *hsr*ω orthologue examined. In the five species (*D. eug, D. moj, D. nas, D. obs* and *D. rho*), for which a contiguous stretch encompassing the conserved splice junctions and the *mir-4951* could not be assembled, the up-stream and down-stream genes were searched in reference to the two scaffolds carrying the *omega* splice junction and *mir-4951*, respectively. Data in Table 2 show the annotated orthologue and the order of some of the up- and down-stream genes that were found to flank the putative *hsr*ω sequences in the examined species genomes. Order of the genes upstream of the *hsr*ω is identical with that in *D. mel* in 29 species and that of downstream genes in 31 species. In the case of *D. ana*, the order of the downstream *mod* and *tin* genes is reversed (Tables 2 and 3). The flanking genes could not be identified in relation to the scaffolds retrieved for *D. ame, D. bus*, and *D. nas*. A similar set of genes flanking the retrieved genomic sequences in most of the examined species further confirms that these scaffolds include *hsr*ω orthologues in other species. We think our inability to identify the flanking genes in *D. ame, D. bus* and *D. nas* (Tables 2 and 3) is most likely due to incomplete genomic data for these cases.

**Table 2.**
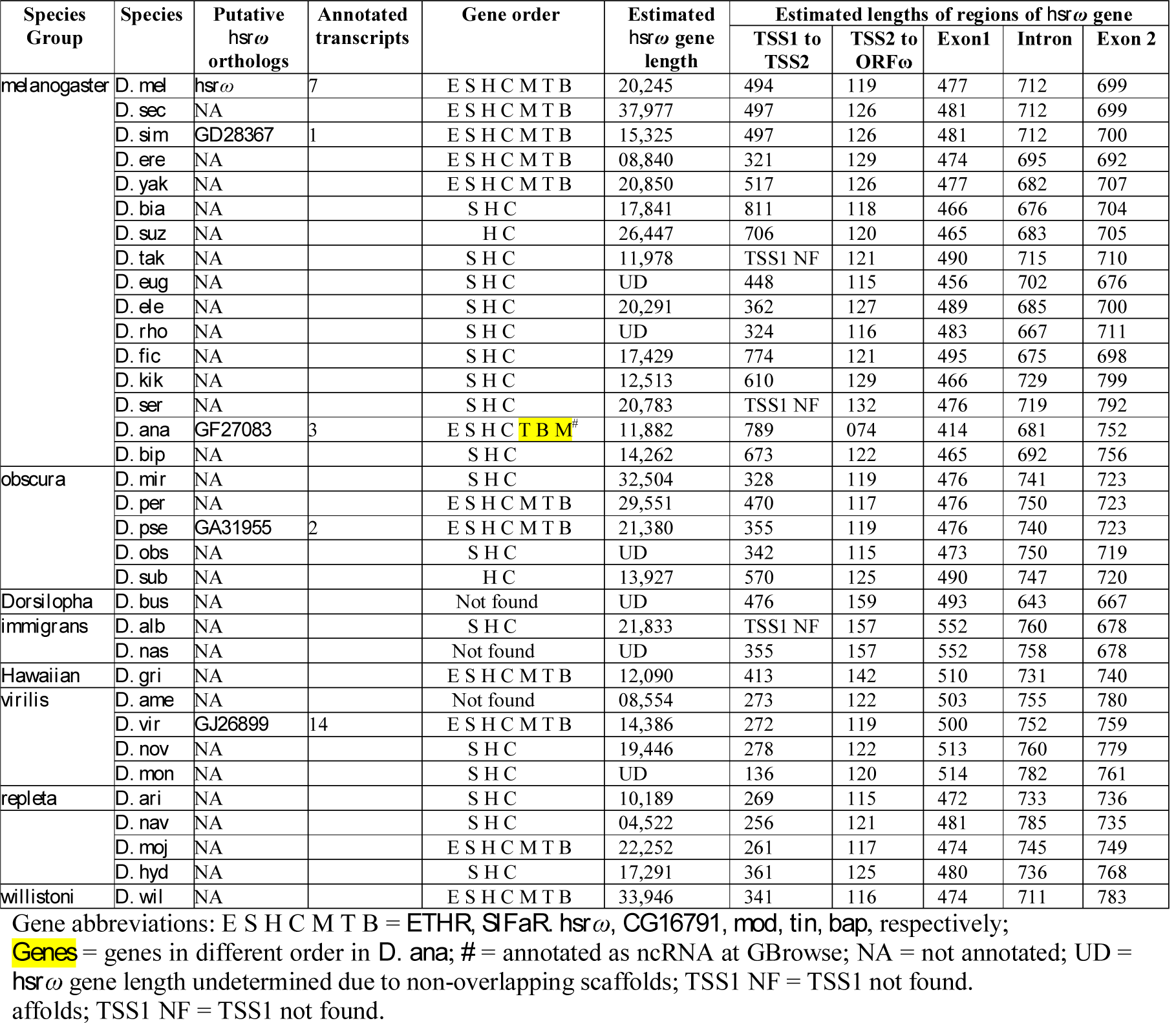
*hsr*ω orthologs, flanking neighbours and lengths of different regions of *hsr*ω gene ascertained on bases of data at http://fb2017_05.flybase.org/cgi-bin/gbrowse/dmel/.

**Table 3.**
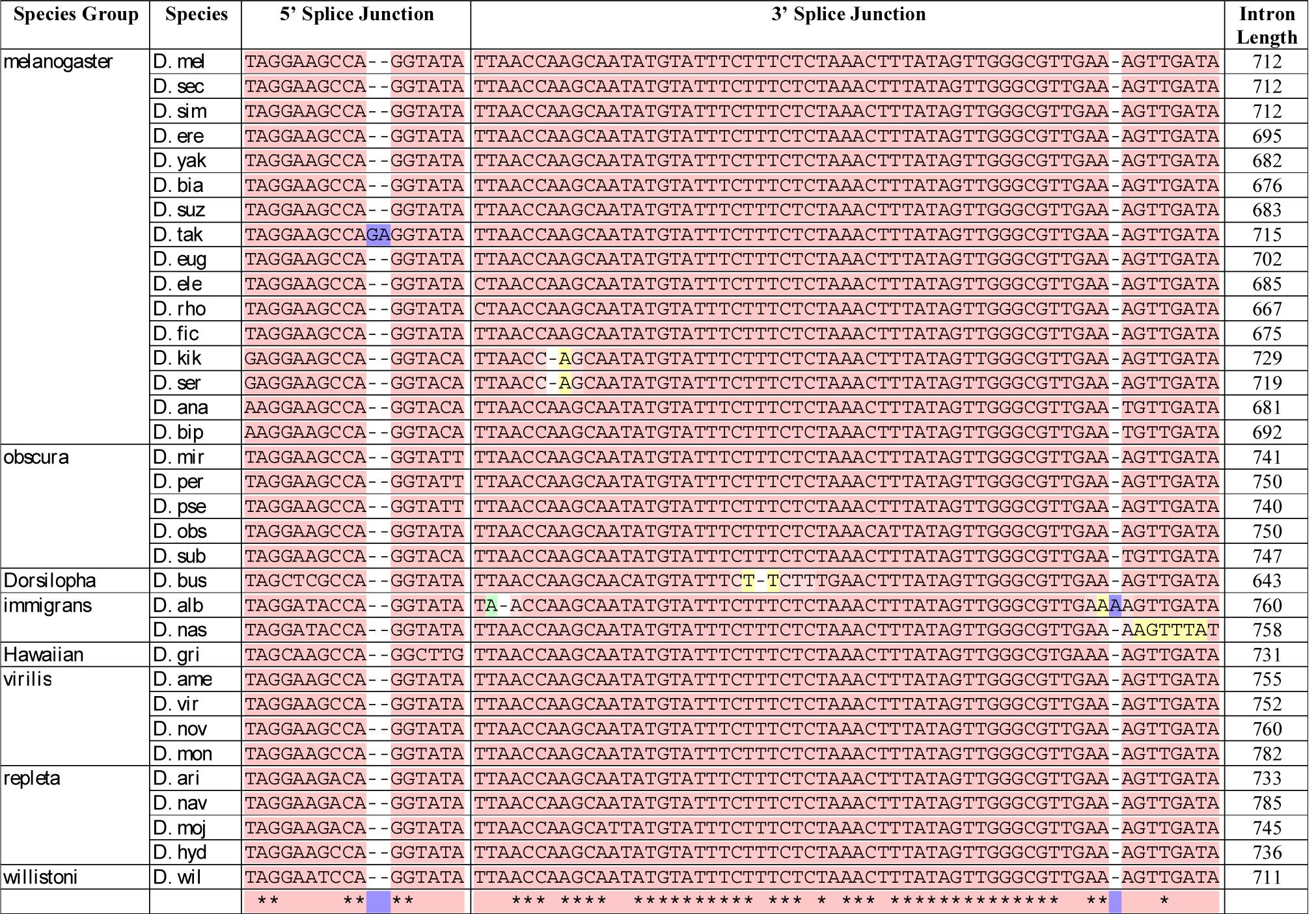
MSA and conservation scores of 16 nt and 60 nt sequences flanking, respectively, the 5’ and 3’ exon-intron splice junctions of the *omega* intron in different *Drosophila* species.

**Fig 3.**
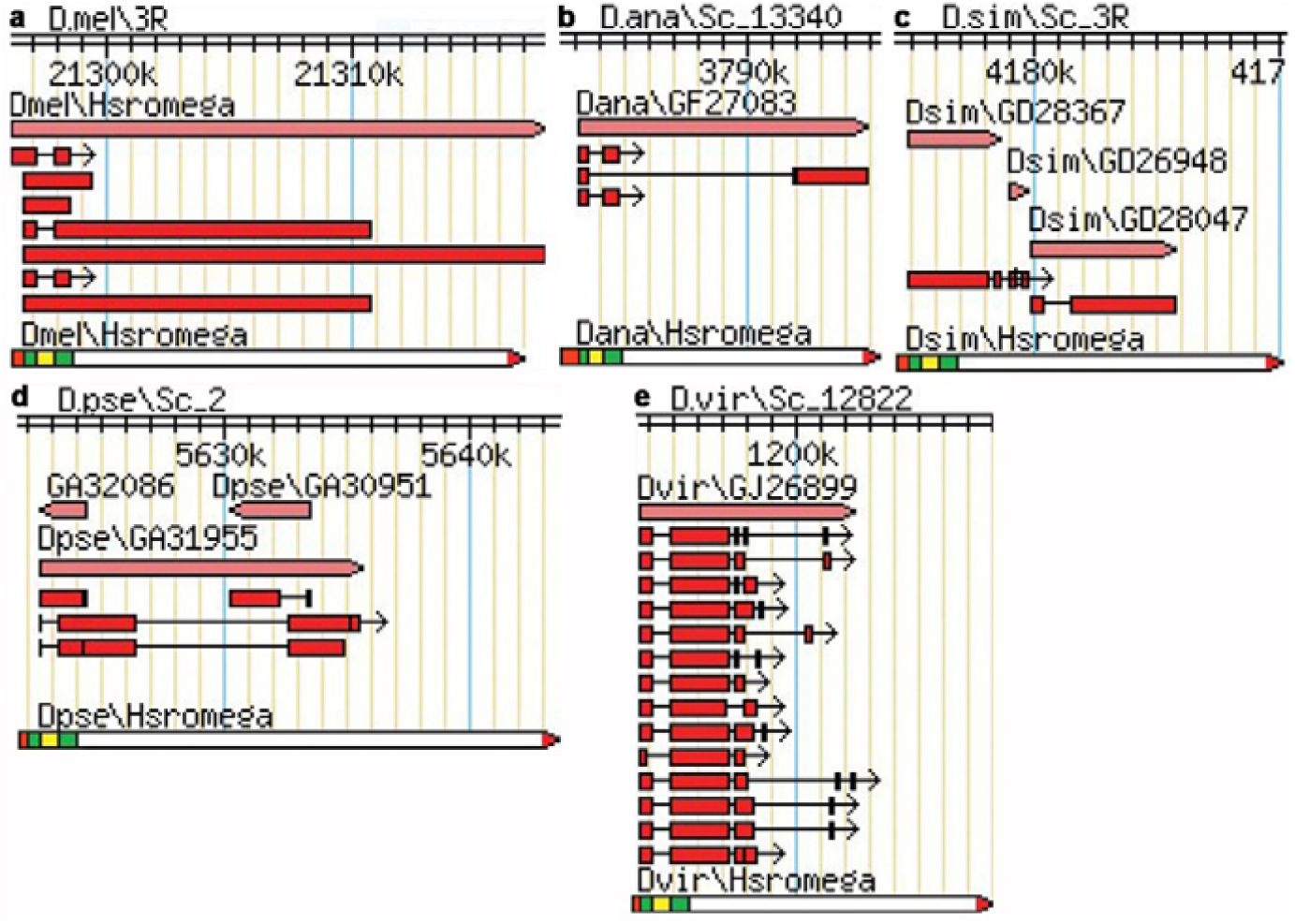
GBrowse genomic co-ordinates of *hsr*ω locus in the 5 annotated *Drosophila* species (*D. mel, D. ana. D. som, D. pse* and *D. vir*) genomes available at the FlyBase. **a-e** Species and the respective contig names are noted above the chromosomal scale for each species. Light red bars denote the *hsr*ω gene with the direction of transcription indicated by the pointed end; the deep red bars represent the annotated transcripts. A manually drawn map at the lower end of each panel denotes the putative *hsr*ω gene, starting with TSS1 and ending at the *mir-4951* as suggested by the present study; the light red box at proximal end corresponds to region between TSS1 and TSS2 (as in *D. mel*), green boxes represent the two exons, the yellow box the *omega* intron while the red arrow head represents the *mir-4951* sequence.

This analysis also clarified a few other issues about the genes flanking the annotated *hsr*ω gene in the 4 species at GBrowse. Two additional ncRNA genes are listed downstream of TSS2 in *D. sim* in the same direction as *hsr*ω. Based on sequence homology revealed by our pair-wise alignment (data not presented), these additional - ncRNA genes appear to correspond to parts of the *hsr*ω gene. Two additional ncRNA genes were found to overlap the *hsr*ω locus in opposite direction in *D. pse* (Fig 3d). The GBrowse data in the current version (FB2020_01) also shows same arrangement of flanking genes in *D. pse* and *D. sim*. It remains to be seen if such instances reflect some errors in sequence/assembly. In *D. sec* genome, an orthologue of *mod* is not annotated. However, we found the *GD15087* gene *D. sec*, located at the corresponding location, to carry a long ORF (including the intronic region) with 94% sequence similarity with the *mod* gene of *D. mel* (data not shown).Therefore, the *GD15087* gene of *D. sec* appears to be an orthologue of the *mod* gene of *D. mel*.

### Sequences flanking the 5’ and 3’ splice junctions in the hsrω gene are ultra-conserved

The MSA analysis revealed that the 16 bp sequence flanking the 5’ splice junction and the 60 bp sequence flanking the 3’ splice junction flanking a similar sized intron in the proximal region of *hsr*ω are very strongly conserved in all the 34 species examined, with conservation scores of 996 and 985, respectively (Table 3). Length of the intron flanked by these exon-intron junction sequences varied between 643-785 bp, generally agreeing with the 712 bp length in *D. mel hsr*ω gene (Table 3). Compared to the highly conserved 5’ and 3’ splice junction flanking sequences, the intron sequences in the 34 species are less conserved, with a score of 549.

A search for splice-sites in the identified *hsr*ω orthologues revealed additional putative splice junctions and thus intronic sequences. However, none of these additional introns were found to be flanked by the 16 bp 5’ and/or 60 bp 3’ splice junctions. In view of the unique features of the single intron in proximal region of the *D. mel hsr*ω gene and its unique conservation across the *Drosophila* species, we designate this as the omega intron.

A search for the ultra-conserved 5’ or the 3’ *omega* splice junction sequences in rest of the genome revealed that their variants were individually detectable in a few other genes in *D. mel*. Eighty two genes (Supplementary Table S1) showed presence of only the 5’ splice junction sequence with 69% to 87% similarity. Interestingly, 41 of these carried the 5’ junction sequence in anti-sense orientation while in the other 41 genes it was in sense orientation (Supplementary Table S1). Only 2 genes, different from those sharing the 16 bp *omega* splice junction sequence, showed 33-36% similarity with the 60 bp *hsr*ω 3’ splice junction. A survey of known/predicted functions of these genes at the FlyBase indicated them to be involved in diverse functions without any apparent commonality in their functions or known interactions (not shown). It is important to note that the 5’ and 3’ splice junctions of the *omega* intron did not occur together in any other gene in any *Drosophila* species.

### The mir-4951 located upstream of the last annotated TTS of D. mel hsrω locus shows conserved presence and location

The *hsr*ω locus in *D. mel* genome is annotated at the FlyBase to include a putative miRNA gene, *mir-4951*, located ∼971 bp upstream of the annotated TTS4, which is at end of the largest transcript *RF* (Fig. 1). In 27 other examined species, the *mir-4951* sequence was present downstream of the *omega* intron on a single contig. Since the conserved *omega* intron and the *mir-4951* are present within <40kb in 28 *Drosophila* species, we believe that the separate contigs identified in *D. eug, D. mon, D. nas, D. obs* and *D. rho* include the proximal and distal parts, respectively, of the *hsr*ω gene. This is further supported by our above noted finding that the genes upstream and downstream of the *omega* intron and *mir-4951* homologous sequence, respectively, were same in 31 examined species.

Therefore, keeping the *D. mel hsr*ω as model, we suggest that the 3’ boundary of the *hsr*ω locus in other *Drosophila* species is close to the 3’ end of *mir-4951* sequence (Figs. 2, 3 and 4). It is expected that further refinements in completion and curation of the genomic sequences in these species would provide more definitive information.

**Fig. 4.**
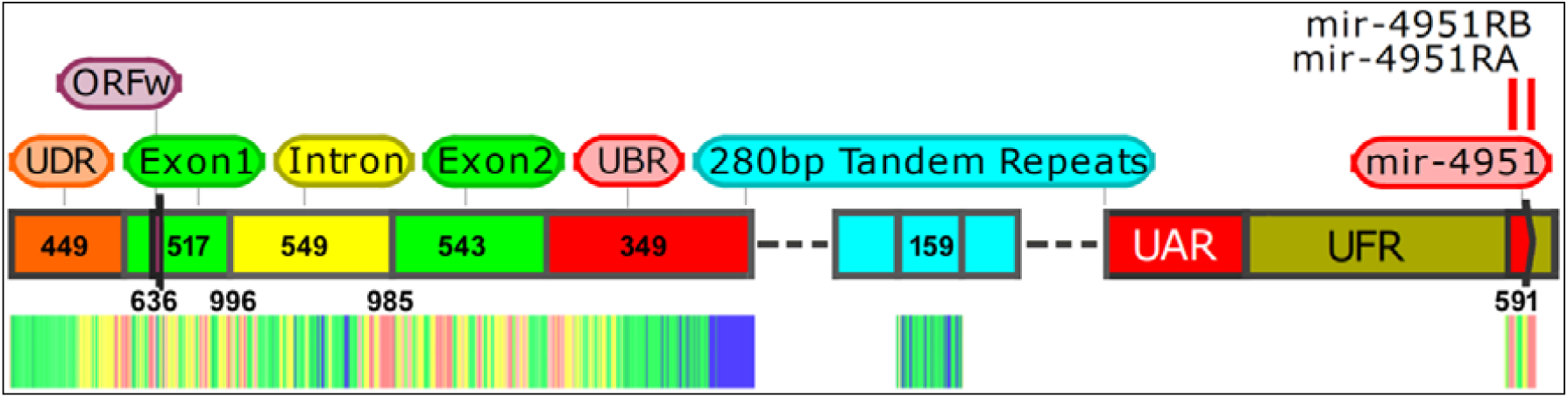
Common architecture but variable sequence conservation across the *hsr*ω gene in different *Drosophila* species. Overall architecture of the *hsr*ω gene in *D. mel* (upper part) and heat map of conservation of different regions taking all the 34 species together (lower part). Numeric values represent conservation score of the corresponding regions.

In the case of *D. bus*, sequence for the full *mir-4951* was not recovered in the entire genome sequence, including the 26 Mb long CP012526.1 *D. bus* chromosome 3R scaffold, which harboured the *omega* intron sequence. Surprisingly, a 45 bp region, located ∼24Mb upstream of the *omega* intron in the *D. bus* chromosome 3R scaffold CP012526.1 showed similarity with the 3’ region of *mir-4951*. It is not clear if this disparity is real or due to some inaccuracies in sequence assembly.

The *D. mel mir-4951* gene is 150 bp long. In other species its length ranges between 131-160 bp. The MSA analysis in 33 species revealed a good conservation with a score of 591, with two regions of the gene showing higher conservation (Table 4). The first region (21 bp), corresponding to the processed 5’ *mir-4951* of *D. mel*, showed high conservation score of 991 in 31 species but appeared more divergent in *D. nav* and *D. nov*. The second region (23 bp) corresponding to the processed 3’ *mir-4951* had a conservation score of 897 in 32 species (Table 4), being more divergent in *D. mon*. These two regions represent the functional mature miRNAs from the locus.

**Table 4.**
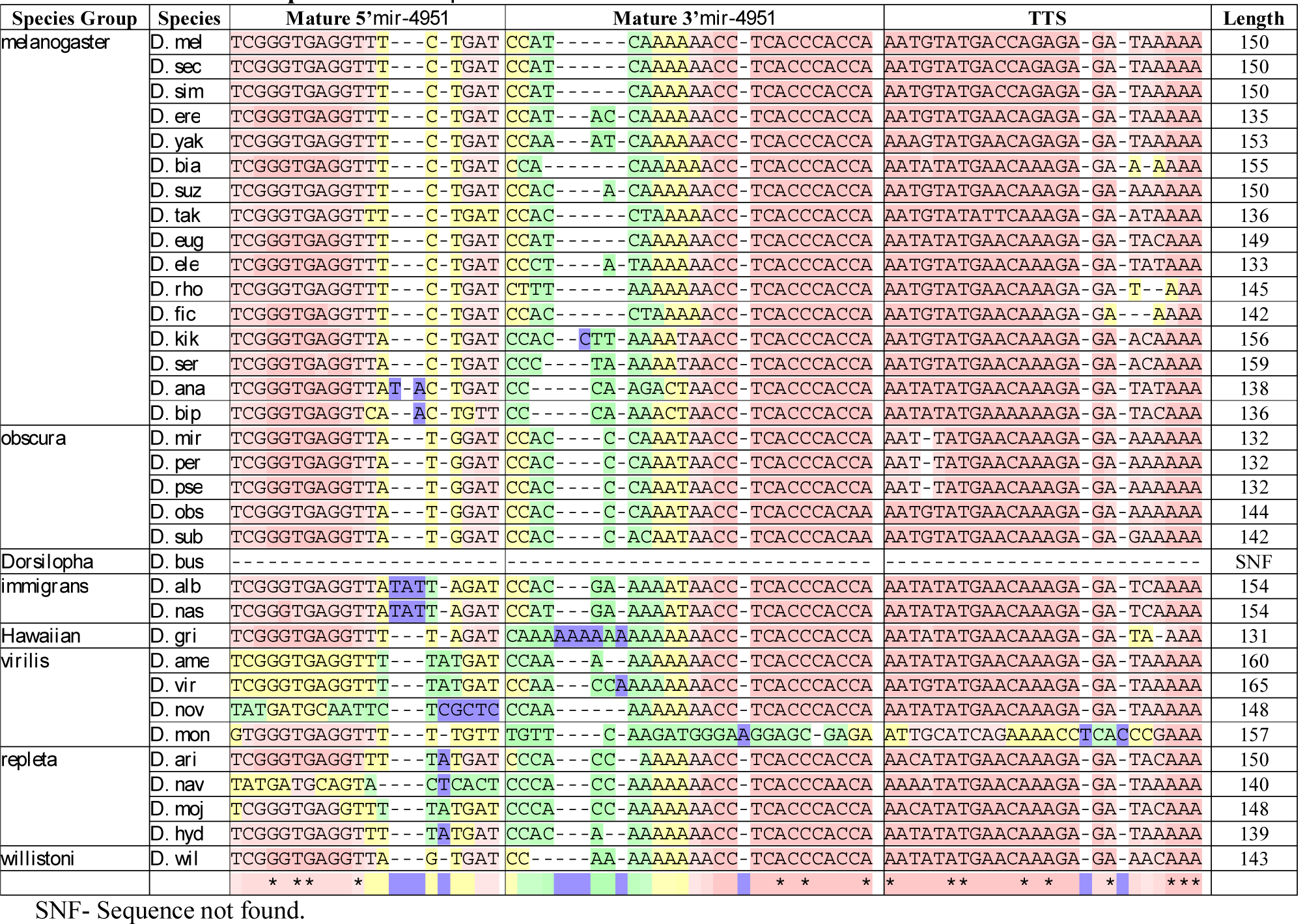
MSA and conservation scores of 5’and 3’ regions of mature *mir-4951* transcripts overlapping with *hsr*ω in different species of *Drosophila*.

### An orthologue of translatable short ORFω is present in all Drosophila species

Earlier studies (Garbe *et al*., 1986; Garbe *et al*., 1989; Ryseck *et al*., 1985) showed that the 1.2kb spliced transcript *RA* or *hsr*ω-c derived from the proximal region of the *hsr*ω gene was cytoplasmic in *D. mel, D. hyd*. and *D. pse*, and that one of the short ORFs, the ORFω, in this transcript was translatable (Fini *et al*., 1989). Therefore, we searched for potential ORFs in proximal part of the *hsr*ω gene. This analysis suggested presence of several putative ORFs, a total of 11 in the entire gene, including 5 ORFs (≥20aa) within the 1.2kb cytoplasmic transcript in *D mel* (Supplementary Table S2).

MSA of the different ORFs identified by the above search using T-Coffee revealed that only the first predicted ORF, located upstream of the 5’ splice junction in each of the 34 species, showed significant degree of conservation with the ORFω of *D. mel* (Table 5). Its location with respect to the TSS2 (see below) and the 5’ splice junction was also comparable in different species. The other putative ORFs in the proximal region (till the end of exon 2 in *D. mel*) of the *hsr*ω gene in different species showed high variability in their amino acid sequence and locations; several longer ORFs (>50aa) were also found in the intron and a few in exon1 and exon2 of the identified *hsr*ω gene region in each of the 33 other species; however, their locations and/or sequences did not show any conservation between the species (Supplementary Table S2).

**Table 5.**
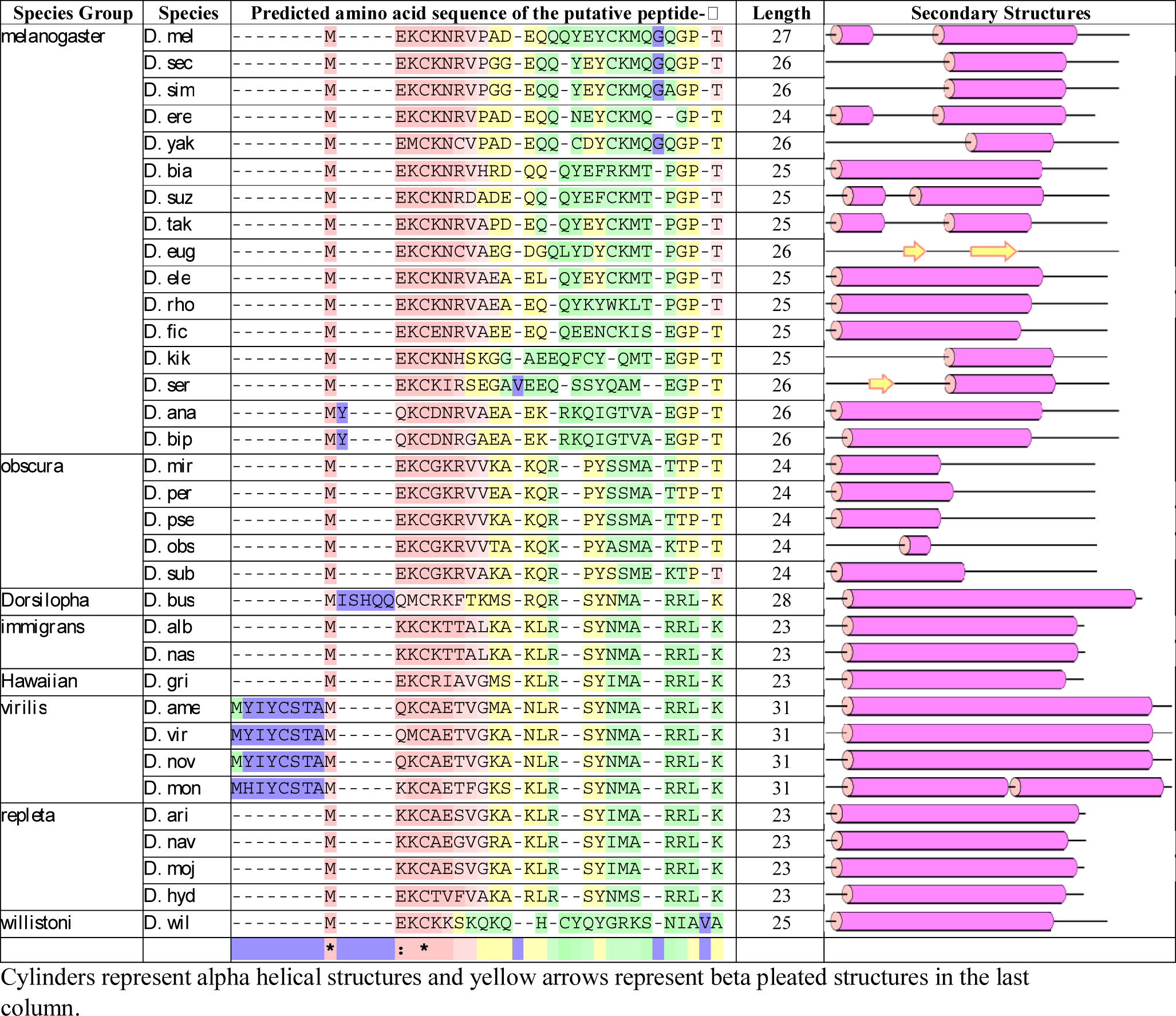
MSA and conservation scores of predicted amino acid sequences of the peptide-▭.

Predicted amino acid sequence of the putative peptide-ω encoded by the ORFω in 34 species (Table 5) showed a conservation score of 627 (Fig. 4). Amino acids in its terminal regions appeared more conserved than those in the middle. The first four N-terminal residues in 27 species are MEKC, MKKC, MQKC, MQMC or MEMC (Table 5). The predicted amino acid sequences of the peptide-ω in 4 species (*ame, mon, nov* and *vir*, members of the *virilis* group) included a few extra residues prior to the MEKC/MKKC/MQKC/MQMC/MEMC first tetrad motif in the other 27 species. In 3 other species (*ana, bip*, and *bus*), extra residues were seen after the first M of the above tetrad. Interestingly, if these extra residues are ignored, the first four N-terminal residues in these seven species would also be similar to those in the other 27 species (Table 5). Therefore, these additional amino acid residues in the predicted peptide-ω in the seven species need further verification of the genomic sequences.

The C-terminal amino acid sequence is “G/TPT” in 21 species and “RRLK” in 12 species, but the peptide-ω ends with “AVA” in *D. wil* (Table 5).

Secondary structure prediction of the putative peptide-ω in different species revealed that a single alpha helix covers >80% of the peptide in 18 species while a shorter alpha helix, covering <50% of the peptide, is seen in 10 species. Two helices covering >70% of the peptide are predicted in 5 species (Table 5). The peptide-ω in *D. ser* had a short alpha helix and a beta pleated structure while in *D. eug* only two beta pleated structures were predicted (Table 5).

### Greater conservation of promoter at TSS2 than at TSS1 and possibility of a novel promoter at ORFω

Initial studies (Garbe *et al*., 1986; Garbe *et al*., 1989; Peters *et al*., 1984; Ryseck *et al*., 1985) on transcripts of *hsr*ω gene identified only one transcription start site (TSS) for the more than one *hsr*ω primary transcripts in *D. mel, D. hyd* and *D. pse*. However, recent annotations at the FlyBase suggest two TSS for *D. mel*. Therefore, we examined the proximal region (∼2kb upstream of the identified 5’ *omega* splice junction) of the retrieved *hsr*ω orthologues in the 33 species for presence of promoter and TSS, in sense orientation, using the BDGP promoter search program.

The TSS1 in *D. mel* corresponds to the site from where the *RD* transcript is initiated (Fig. 1). Promoter search analysis suggested presence of a promoterat the corresponding site in 31 species. However, a promoter was not predicted at a corresponding region in *D. alb, D. tak* and *D. ser*, The predicted TSS1 promoter showed a low degree of sequence conservation between the 31 species (Table 6A). The region between TSS1 and beginning of exon-1 (beginning at the TSS2, see below) in *D. mel* was annotated as UDR (Unique to RD transcript region; Fig. 1); lengths of its putative orthologue in other species ranged from 136-811 bp (Table 2).

**Table 6.**
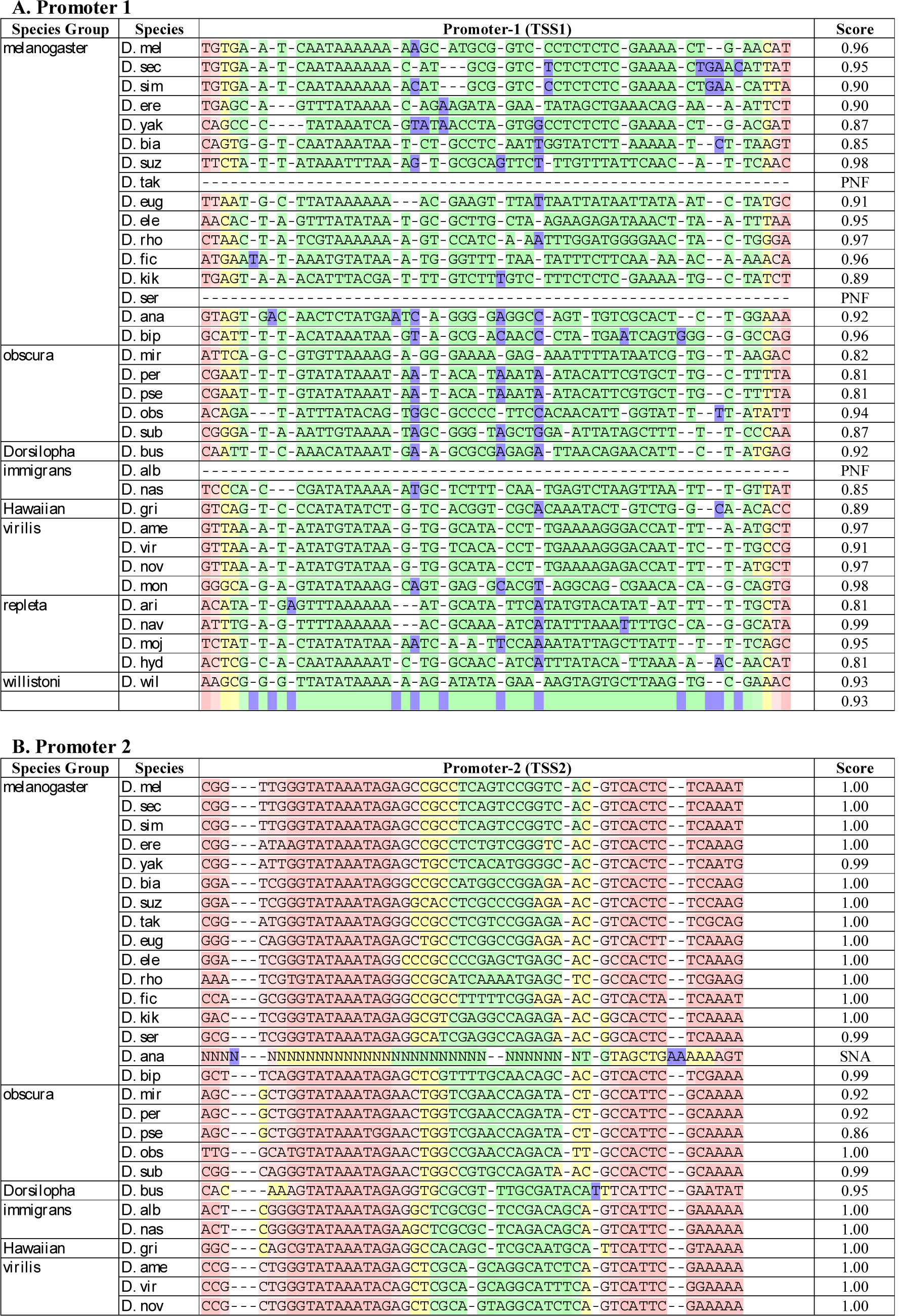
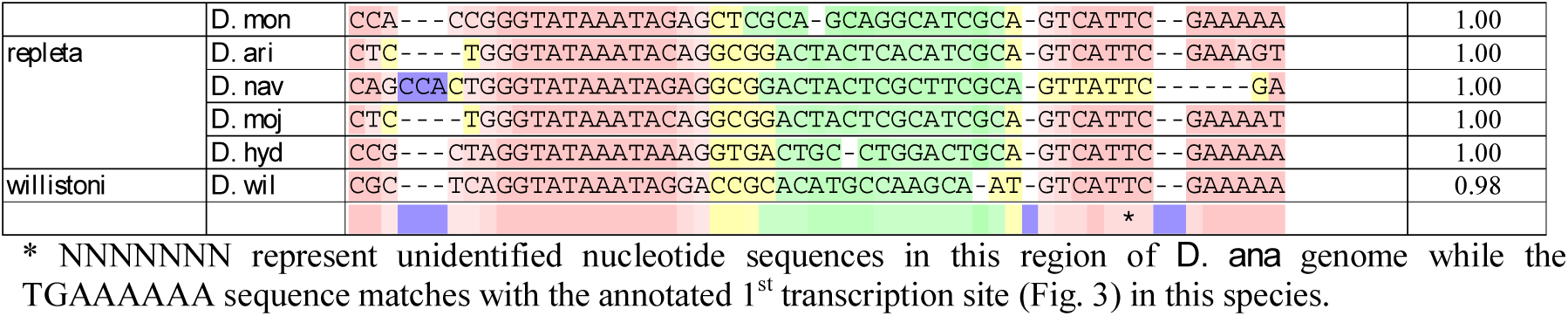
Predicted promoters 1 and 2 of *hsr*ω in different species of *Drosophila*.

Search for homology for the TSS2 of *D. mel* revealed presence of a promoter at ∼115-130 bp upstream of the ORFω in 31 species and at ∼160 bp upstream in 3 other species (Table 2). MSA of the predicted TSS2 promoters of the 33 species revealed a very high homology score with that of *D. mel*, ranging from 0.86-1.00 in different species (Table 6B). The ambiguous sequence available for this region in *D. ana* (81Ns noted in the shotgun sequence scaffold, http://flybase.org/cgi-bin/gbrowse2/dana/scaffold_13340:3,782,971...3,783,051), precluded a definitive identification of the TSS2 promoter in this species. However, the GBrowse transcript annotation data (Fig. 3b) suggests presence of a functional promoter at 74 bp upstream of the ORFω in this species, which partially matched with the putative TSS2 identified in other species (Table 6B).

A 14 bp region (GGTATAAATAGAGC) located 18 bp upstream of the annotated TSS2 in *D. mel* is highly conserved and based on its sequence, this may function as the TATA box for TSS2. The 477 bp region starting from TSS2 and ending at the 5’ splice junction is annotated as exon 1 in *D. mel*, and correspondingly, the region between the predicted TSS2 and the 5’ splice junction is named as exon 1 in the other 33 species, with its length varying from 414 to 552 bp. Although exon-1 as a whole showed only moderate conservation with a score of 517, a 45 bp region at the 5’ end of exon1 showed high conservation; another stretch of 37 bp, located 140 bp downstream of the ORFω in *D. mel* exon-1 also showed good conservation in different species (Fig. 4).

A comparison of the relative locations of the annotated and predicted TSS1 and TSS2 in the four other (*D*.*ana, D. sim, D. pse* and *D. vir*) species whose genome annotation data are available at the FlyBase revealed that in most cases the coordinates of predicted TSS2 matched with those of the annotated transcript sites. As noted above, the TSS1 is annotated only in the case of *D. mel hsr*ω, which fully matched with the predicted TSS1 I in this species. The TSS2 is annotated for all the 4 other species and interestingly, in three of them the predicted TSS2 nearly matched with the respective annotated TSS. In the case of *D. ana*, this match could not be ascertained because of ambiguity in the available sequence for this region. In the case of *D. pse*, the *hsr*ω gene region annotation at Flybase suggests transcription to start downstream of the predicted TSS2 (Fig. 3). However, in view of the high similarity between the predicted TSS2 in *D. pse* with that in other species, we believe that further curation of the *hsr*ω transcripts in these species is needed.

We suggest that the region between TSS1 and 5’ splice junction of the *omega* intron in *D. mel* be named as exon 1a since its 3’ end is same as that of the exon 1. It, however, remains to be seen if a functional TSS1 and a transcript corresponding to *D. mel hsr*ω-*RD* is indeed present in other species.

Our promoter search revealed a new potential transcription start site, which has so far not been annotated in *D. mel* or any other species. The core promoter of 50 bp for this potential third TSS (TSS3), predicted by the BDGP promoter search tool, overlaps with the ORFω (Supplementary Table S3) and is present in 23 species with a score of 0.82-0.99. This predicted promoter sequence shows moderate conservation with a score of 432 (Supplementary Table S3). It may be noted that although a TSS has not been annotated at this region in *D. mel* or any other species at the FlyBase, two cDNA sequences (BT122054.1 and M13240.1) in the NCBI database (Garbe & Pardue, 1986) start 93 nt and 26 nt, respectively, upstream of this predicted promoter (Fig. 1).

It is notable that annotations for the *hsr*ω orthologues in *D. pse* and *D. sim* at the Fybase indicate additional transcripts that start far downstream of the TSS2 (Fig. 3). In addition, the Flybase annotation indicates two anti- sense transcripts of *hsr*ω gene in *D. pse* (Fig. 3).

### First transcription termination site and exon 2 regions are variably conserved

The FlyBase has annotated 4 transcription termination sites (TTS) on the *hsr*ω gene of *D. mel*. The first TTS (TTS1) is 699 bp downstream of the 3’ splice site of the *omega* intron, and this region annotated as exon 2 in *D. mel*. The PolyAh output suggested that a TTS is present between 667 to 799 bp downstream of the 3’ splice junction in the *hsr*ω gene in all the 34 species. In keeping with the annotation in *D. mel*, we named this TTS as TTS1 and the region between the *omega* intron’s 3’ splice site to TTS1 as exon 2 (Fig. 4). However, the predicted TTS1 matched with the known TTS at the end of exon 2 only in *D. mel*. In other species for which transcripts are indicated at the FlyBase (Fig. 3), the locations of predicted TTS1 and the annotated TTS1 did not match well. Nevertheless, we considered the sequence between 3’ splice site of the *omega* intron and the predicted TTS1 as exon 2 in all the species. Despite the comparable length of exon 2 in different species, its overall conservation score was relatively poor with a score of 543, although some regions along its length showed a higher degree of conservation. These included 75 bp and 30 bp stretches, 199 bp and 379 bp, respectively, downstream of the 3’ splice junction in *D. mel hsr*ω gene. A 147 bp stretch, 458 bp downstream of the 3’ splice site, also showed a moderate conservation in different species (Fig. 4).

### Multiple transcription termination sites may exist in hsrω orthologs in other species

We used PolyAh and other tools like ARNold (http://rna.igmors.u-psud.fr/toolbox/arnold/index.php), PolyA Signal Miner (http://dnafsminer.bic.nus.edu.sg/PolyA.html), POLYAR (http://cub.comsats.edu.pk:8080/polyar.htm) PASIGNAL (http://cub.comsats.edu.pk:8080/GEFWebServer.htm) to identify other potential TTS. The PolyAh analysis predicted a novel TTS at a distance of 20-27 bp (Table S4) upstream of the TTS1 in all species except *D. wil*. None of these TTS prediction tools definitively predicted other TTS sites, including those that are annotated for the four species at the FlyBase (Fig. 3). Search for other TTS using the PolyAh tool generated very large numbers of potential termination sites in all the species, with little conservation in terms of their locations. The only other TTS consistently predicted by the PolyAh tool was located downstream of the *mir-4951* in 17 species (*D. ana, D. bip, D. ere, D. gri, D. mel, D. mir, D. moj, D. obs, D. per, D. pse, D. rho, D. sec, D. ser, D. sim, D. sub, D. suz and D. tak*). In the other species, a TTS could not be definitively identified downstream of the identified *mir-4951* sequence.

### Tandem repeats: the most diverse region of hsrω gene

A major part of the *D. mel hsr*ω gene contains tandem repeats of 278 bp (generally denoted as 280 bp repeat), unique to this gene and which typically account for ∼50% of the 21.7kb long gene (Table 7). These are located between the TTS2 and TTS3 of the *D. mel hsr*ω (Figs. 1 and 4).

**Table 7.** Repeat units in the *hsr*ω *genes* in different species of *Drosophila*.

| Species Group | Species | Lengths of repeat units and their tandem repetition frequencies | Number of ATAGGTAGG Nonamer (and variants) # | Total Estimated Length |
| --- | --- | --- | --- | --- |
| *melanogaster* | *D. mel* | 87x3, 278x37, 365x2 | 68 | 10,535 |
|  | *D. sec* | 76x8, 280x86, 354x11.5 | 129 | 26,788 |
|  | *D. sim* | 278x12 | 29 | 03,331 |
|  | *D. ere* | 114x2, 230x5.5 | 12 | 00,481 |
|  | *D. yak* | 89x56, 128x2.2, 180x14.5, 220x4, 288x36, 376x2, 463x9.5 | 31 | 09,349 |
|  | *D. bia* | 82x2, 152x1.8, 180x2, 228x16, 254x7, 338x2, 379x2, 483x2.5 | 36 | 14,912 |
|  | *D. suz* | 128x2.1, 310x2.5, 336x4.2, 354x5 | 41 | 04,306 |
|  | *D. tak* | 204x4, 242x7, 490x2 | 18 | 03,396 |
|  | *D. eug\** | 61x14, 78x4, 87x2, 96x39, 157x16, 175x5, 195x2.5, 253x6 | 03 (+14 TGTGCTAGG) | 04,542 |
|  | *D. ele* | 55x2, 164x2, 180x2, 232x3 | 01 (+27 TTAGGTAGG) | 01,010 |
|  | *D. rho\** | 142x2, 260x11.6 | 02 (+28 TTAGGTAGG) | 03,027 |
|  | *D. fic* | 77x2 | 11 | 00,145 |
|  | *D. kik* | 62x18.6, 176x17, 245x2.2, 309x4.2, 350x8.4, 429x2.2 | 31 | 04,144 |
|  | *D. ser* | 67x33, 160x25, 213x14, 232x2, 280x15, 368x2, 468x3, 493x10.3 | 53 | 09,156 |
|  | *D. ana* | No repeats found | 04 (+14 TTAGGTAGG) | 00,000 |
|  | *D. bip* | 179x3.5 (located downstream of *mir-4951*) | 03 (+16 TTAGGTAGG) | 00,636 |
| *obscura* | *D. mir* | 343x2, 356x3.7 | 27 | 02,167 |
|  | *D. per* | 156x2, 354x9 | 43 | 03,604 |
|  | *D. pse* | No repeats found | 22 | 00,000 |
|  | *D. obs\** | No repeats found | 05 | 00,000 |
|  | *D. sub* | No repeats found | 01 | 00,000 |
| *Dorsilopha* | *D. bus* | 61x2, 135x5.4 | 06 | 00,722 |
| *immigrans* | *D. alb* | No repeats found | 20 (+12 TTAGGTAGG) | 00,000 |
|  | *D. nas\** | Sequence not available | 00 | ----- |
| *Hawaiian* | *D. gri* | 60x5, 110x2, 155x25.7, 464x8.6 | 01 (+27 CTAGGTAGG) | 04,205 |
| *virilis* | *D. ame* | No repeats found | 00 (+02 TTAGGTAGG) | 00,000 |
|  | *D. vir* | 36x236, 83x27.5, 112x2, 124x6, 137x41, 220x14.4, 273x2.7, 356x6.2, 437x2 | 25 | 05,883 |
|  | *D. nov* | 85x57.7, 99x2, 140x23, 215x27, 300x2, 365x8, 455x2.4 | 00 (+85 TTAGGTAGG) | 11,078 |
|  | *D. mon\** | 87x3, 132x2.2, 162x4, 223x2.3, 294x2.2 | 00 (+05 CCTATTAGG) | 01,386 |
| *repleta* | *D. ari* | No repeats found | 00 | 00,000 |
|  | *D. nav* | No repeats found | 00 | 00,000 |
|  | *D. moj* | 60x187, 124x56.6, 156x1.8, 180x49.5, 246x38.6, 415x1.8, 483x9.8 | 00 (+76 TTAGGTAGG) | 11,935 |
|  | *D. hyd* | 57x2, 115x92, 170x4, 190x2, 228x44, 246x2, 353x2.3 | 45 (+39 TTAGGTAGG) | 10,826 |
| *willistoni* | *D. wil* | 63x2, 135x2, 197x125, 332x2 | 00 (+58 ATAGGCAGG) | 24,363 |

Tandem Repeat Finder tool revealed variable sizes and numbers of repeats downstream of the TTS1 in 25 species. Intriguingly, tandem repeats were not found in the identified *hsr*ω orthologue sequences in 8 species while the absence of genomic sequences for an unknown length of middle part of the identified *hsr*ω gene region limited the search for tandem repeats in the case of *D. nas*. In some species, e.g. *D. per, D. ser, D. rho*, the repeats were not tandemly arranged since some unique sequences intervened between the repeats (Fig. 5). The repeat lengths and the number of copies of tandem repeats in 25 species are shown in Table 7.

**Fig. 5.**
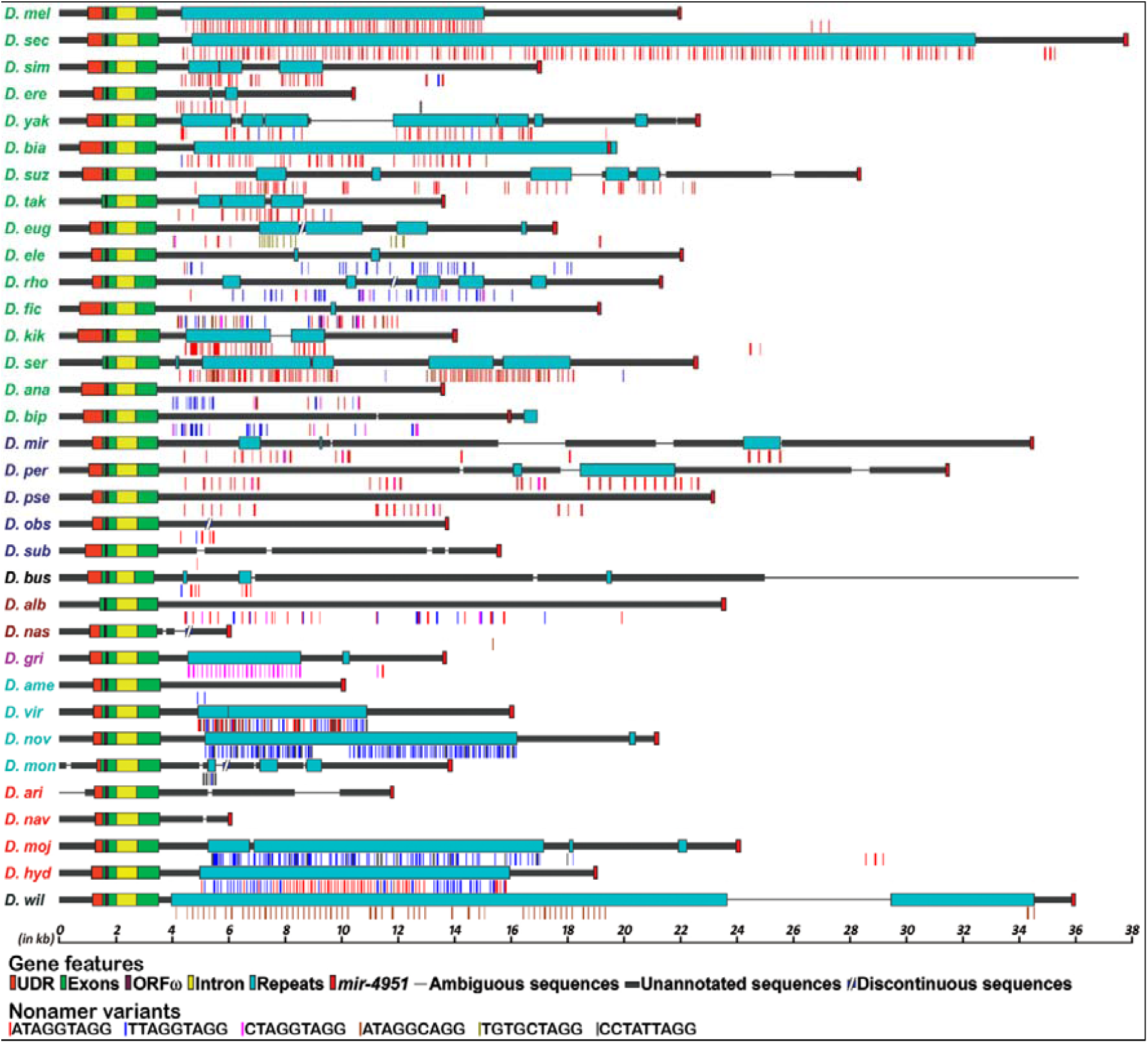
Predicted architecture of *hsr*ω gene in the genus *Drosophila*. A full length SnapGene annotated map of *hsr*ω in all species beginning from 2kb upstream of 5’ splice junction and ending with *mir-4951*. Different domains in this gene includes two exons (green), the ORFω (purple), the *omega* intron and the UDR (orange) in the proximal part and the *mir-4951* at the 3’ end and the most diversified tandem repeat region (sky-blue) in the middle part; thick black horizontal and thin light grey lines in the gene region indicate unannotated sequences with little inter-species conservation and ambiguous sequences, respectively. The discontinuity in six species indicates that the two scaffolds, presumably representing proximal and distal regions of the *hsr*ω, did not show overlap to enable assembly of a contiguous region. The coloured thin vertical lines below each gene map indicate locations of the conserved nonamers with the specific nonamer sequence represented by each colour as noted at the bottom.

It is significant that these repeats were found in the total genome only in the identified *hsr*ω gene region in each species. In agreement with earlier report (Garbe *et al*., 1986), the base sequences of these repeats showed high variations between species, resulting in a very low conservation score (Fig. 4).

A Percent Identity Matrix (PIM) analyses of the different repeat units in each of the 22 species, which had displayed multiple repeat unit sizes, revealed variable degrees of divergence within a species (Fig. 6). The PIM suggests that, most instances of the variable lengths of tandem repeat units in a species are variants of tandem repeats of one or more of smaller repeat units. However, in *D. bia* (Fig. 6e), *D. gri* (Fig. 6p), *D. nov* (Fig. 6r), *D. mon* (Fig. 6s), *D. moj* (Fig. 6t) and *D. wil* (Fig. 6v), one of the *hsr*ω orthologue associated repeat units appeared to be very different from the other units in the same species.

**Fig. 6.**
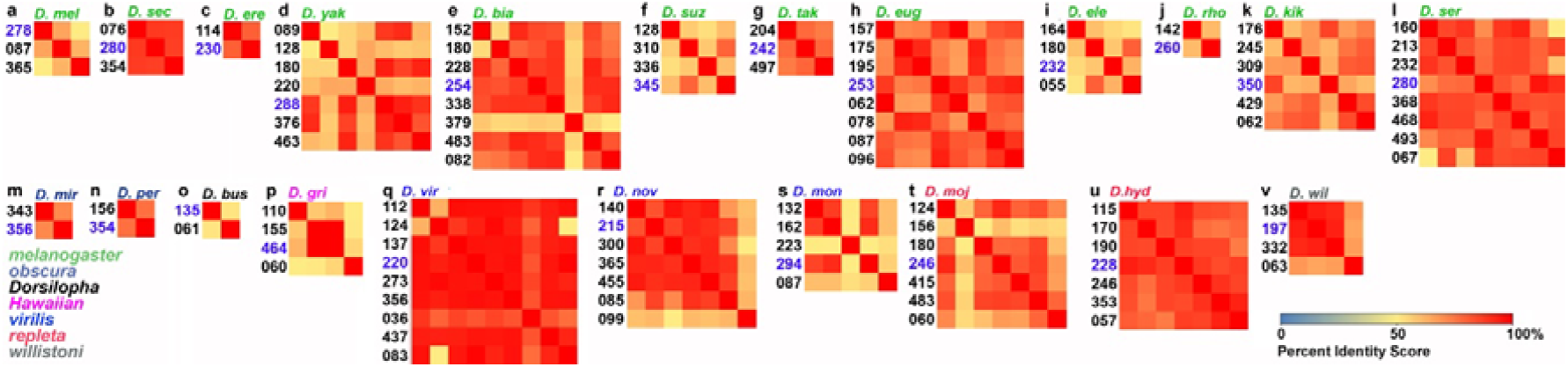
(a-v). Percent Identity Matrix of the tandem repeat units in each of the 22 species with more than one tandem repeat units in the *hsr*ω gene region reveals variable degree of divergence within a species. Lengths (in bp) of the different repeat units in different species (named on top with different colour fonts indicating species groups) are shown on left of each panel. Colour codes for species groups and for heat-maps are given on lower left and right corners, respectively. The repeat units marked in blue fonts in each species were used for PIM analysis of the inter-specific similarity of tandem repeats shown in Fig. 8.

A nonamer sequence (ATAGGTAGG) was earlier reported (Garbe *et al*., 1986) to be repetitively present in the otherwise diversified tandem repeats in *D. mel* and *D. hyd*. Our present results also showed that despite the high variability in tandem repeat sequences within and between species, multiple copies of this classical nonamer and/or its variants (TGTGCTAGG, TTAGGTAGG, CTAGGTAGG, CCTATTAGG, ATAGGCAGG) are present with varying frequencies in 31 species, including in those that do not show tandem repeats (Table 8). As shown in Fig. 5, the nonamers, when present, were mostly restricted to the tandem repeats. Interestingly, even in species which did not show any tandem repeats, the classic and/or variant nonamer sequences were consistently restricted to the region downstream of the TTS1 and upstream of the *mir-4951*, which would correspond to the region where tandem repeats are located in the other species. They were completely absent in the proximal region (TSS1 to TTS1) of the gene in all species (Fig. 5).

A genome wide blast search revealed that in *D. mel* genome, the ATAGGTAGG nonamer is present at several other loci also. The nonamer sequence appears to be more common at centromeric heterochromatin of 3R, rDNA and at telomeres of X and Y. Such motifs are also present at other regions but with a lower than the expected random frequency of a nonamer.

### Very low conservation immediately upstream and downstream of the tandem repeat region

The unique sequence regions immediately upstream (UBR, Unique region upstream of the tandem repeats between TTS1 and TTS2 in *D. mel*) and downstream (UAR, Unique region after the stretch of tandem repeat till the TTS3 in *D. mel*) regions flanking the tandem repeat sequences were also examined for conservation in different species. Since the UBR in *D. mel* is 888 bp long, a similar length of sequences downstream of the TTS1 in each of the 34 species was examined by MSA. This revealed that the proximal ∼50 bp sequence was highly conserved while the middle region was much less conserved and the ∼200 bp at its 3’ end showed no conservation at all (Fig. 4). In contrast, a similar MSA analysis for the UAR sequence of *D. mel* in different species showed its conservation only in the 4 species closely related to *D. mel*. The *D. sec* and *D. sim* showed ∼70% while *D. yak* and *D. ere* showed ∼20% conservation for the UAR region. Little conservation was seen in other species.

MSA of the UFR (Region, unique to the *hsr*ω*-RF* transcript, downstream of TTS3 and upstream of the *mir-4951* in *D. mel*) revealed detectable similarity only in 3 species, viz., *D. sim* (∼80%), *D. sec* (∼70%) and *D. ere* (∼30%). In view of such limited conservation across the 34 species, the conservation heat maps for the UAR and UFR could not be prepared.

### Evolutionary relationship of hsrω gene in different species of Drosophila

Based on the above analyses, a schematic of the *hsr*ω gene, beginning at 2kb upstream of the 5’ splice junction of the *omega* intron and ending with the *mir-4951* gene sequence at the 3’ end, in the 34 examined species of *Drosophila* is presented in Fig. 5. We have used the *D. mel hsr*ω gene architecture as the base, and accordingly, have tentatively modelled the *hsr*ω gene in all species to end with the *mir-4951*. It may, however, be noted that only in the cases of *D. mel* and *D. ana* the longest annotated transcript from this gene covers the *mir-4951* sequence since in the 3 other annotated *Drosophila* species’ genomes available at the FlyBase, the longest annotated *hsr*ω transcripts terminates much before the *mir-4951* sequence (Fig. 3). As noted earlier, the transcription start site annotated at GBrowse for the *D. pse hsr*ω orthologue differs from that predicted by our bioinformatics analysis. We believe that as more exhaustive genome and transcriptome data for different species become available, a better definition of the *hsr*ω gene and its transcripts would be possible. Till that time, we prefer to use the *D. mel hsr*ω gene’s architecture as the model for the *hsr*ω orthologue in all the species (Fig. 5).

In order to find evolutionary relatedness of the *hsr*ω gene in *Drosophila* species, we used sequences from different regions (UDR, Exon-1, Intron, Exon 2, ORFω, UBR, Repeats, and *mir-4951*) to generate phylogenetic trees (Fig. 6). MEGA X software was used to generate the phylogenetic tree following MSA by T-Coffee tool. MEGA X, uses Maximum Likelihood (ML) algorithm Tamura Nei model with nearest neighbour interchange, (Tamura & Nei, 1993) to construct the topology. The species names have been color-coded in Fig. 6 to indicate the 8 sub-genus/species groups to which the different species belong (O’Grady & DeSalle, 2018).

For each analysis, branch support was calculated by bootstrap analysis consisting of 500 replicates to generate phylogenetic trees of different regions (Fig. 7). Except for members of the *melanogaster* species group, those of the other groups tended to be bunched together in the phylogenetic maps for all the regions, except the repeat region (Fig. 7g), which did not reflect any phylogenetic relation across the different species groups. Members of the more diverse *melanogaster* species group were in most cases, however, clustered in two or even three separate groups although each of these clusters included species that are generally considered to be phylogenetically closer. Significantly, the phylogenetic map of the *omega* intron (Fig. 7c), excluding the ultra-conserved splice junction regions, fully matched the reported phylogenetic relationships between different species of *Drosophila* (Bai *et al*., 2007; O’Grady & DeSalle, 2018) since in addition to clustering of the 16 *melanogaster* group species in connected groups, other species also displayed the expected phylogenetic relationship.

**Fig. 7.**
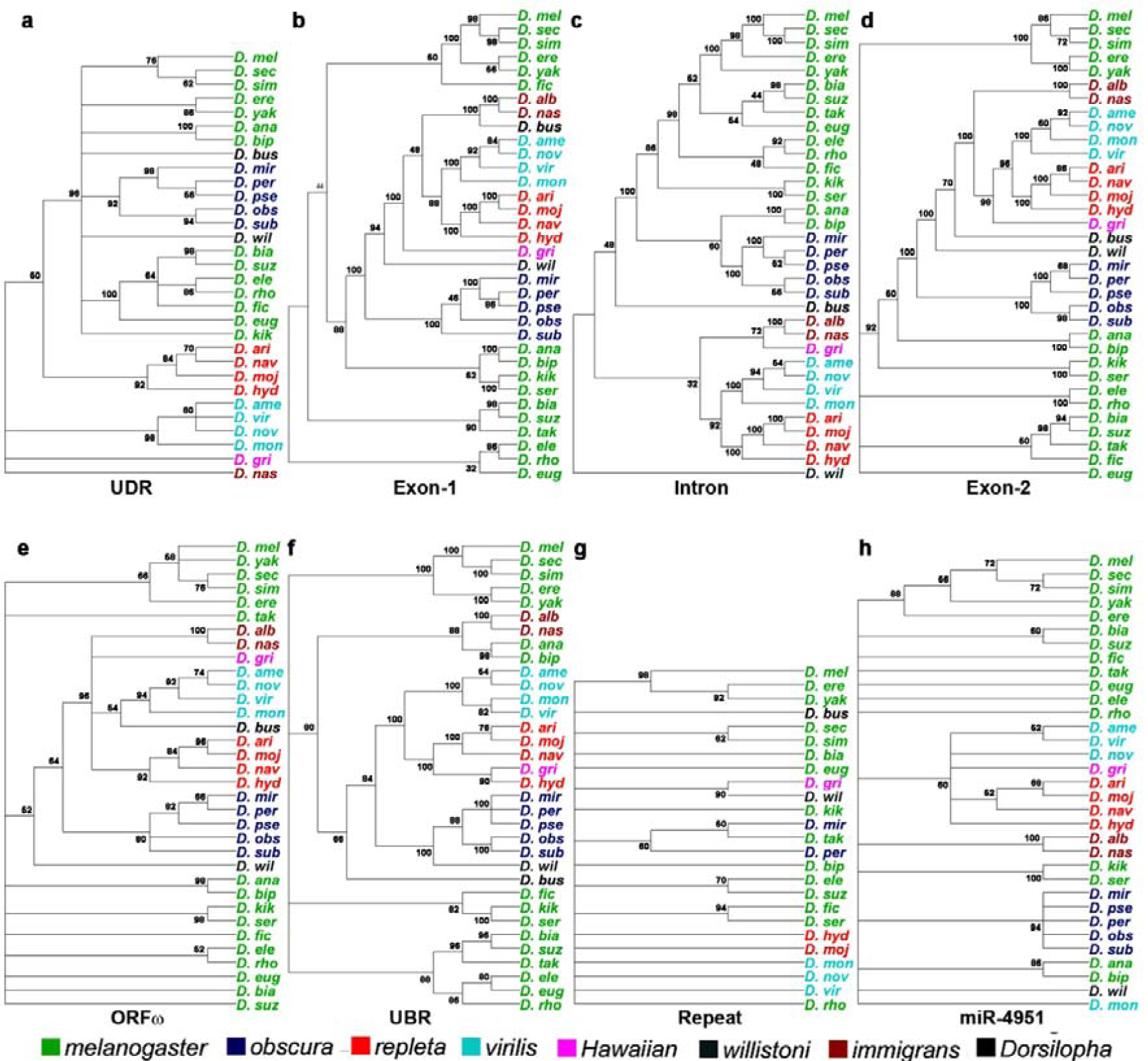
Molecular phylogenetic relationships for different regions of the *hsr*ω gene by Maximum Likelihood method. Phylogenetic trees for different regions (UDR, Exon-1, Intron, Exon-2, ORFω, UBR, Repeat and *mir-4951*) of *hsr*ω gene from 34 species are shown in **a-h**. Numeric value at the base of each branch point represents the homology index between two branches.

We also analysed percent identity matrix (PIM) to examine the divergences at different regions of the *hsr*ω gene in *Drosophila* species (Fig. 8a). The exon-1, *omega* intron, ORFω, exon2 and the *mir-4951* showed higher conservation within species subgroups. The exon-1 and exon-2 regions in the *melanogaster* species group appear to share greater similarity with the *obscura* group than with other species. Within the *melanogaster* species group, *D. ana* and *D. bip*, which belong to *ananassae* sub-group, show greater divergence for all regions from the other species in the *melanogaster* group. As revealed by the phylogenetic tree analysis (Fig. 7), the PIM analysis also showed least conservation of the repeats associated with this gene. Accordingly, a relatively high PIM value for the tandem repeats was detectable only between very closely related species, even within a species sub-group. Blast search for sequence similarity for the repeat units within closely related species (Fig. 8b) revealed that some similarity in the tandem repeat units is detected only between very closely related species since with increasing phylogenetic distance, the similarity becomes increasingly less. For example, within the *melanogaster* species group, *D. mel* tandem repeat unit shares greater similarity only with that in *D. sim* and *D. sec* but hardly with tandem repeats in other species (Fig. 8b). Despite such rapid divergence of the tandem repeats, the nonamer sequences, however, remain abundant in the repeat/distal region of most species (Fig. 5).

**Fig. 8.**
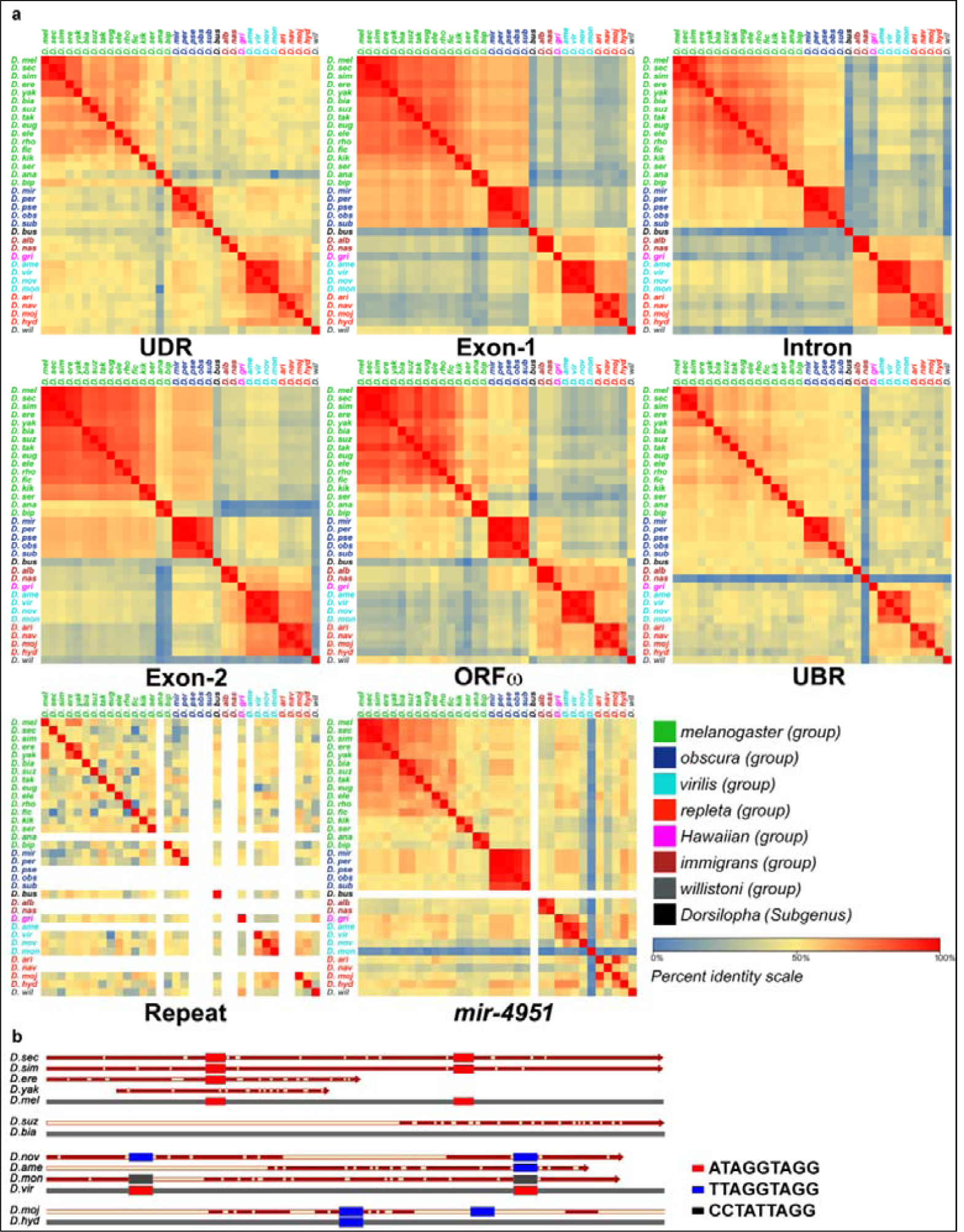
a. Percent Identity Matrix analysis confirms variable but independent divergences in sequences in different domains of the *hsr*ω gene in the 34 Drosophila species. Colour codes for heat-maps and for species groups are given below the matrices. White Rows and columns in PIM for Repeats and *mir-4951* reflect absence of the corresponding sequences in some species. **b**. Comparison of length and sequence of the tandem repeat units in related species in species sub-groups: single repeat units of *D. mel* (278 bp), *D. bia* (254 bp), *D. vir* (220 bp) and *D. hyd* (228 bp), shown as grey thick lines at bottom in each group (not to scale), are compared with the best matching repeat units of related species (shown above the four species units); solid brown regions indicate sequence similarity while white rectangles show sequence mismatches with the master repeat unit; the conserved nonamers are shown as solid rectangles with colour code as in the legend on right;

Phylogenetic trees and the PIM show that the sequences for different regions of the *hsr*ω gene have diversified rather independently. Further, while within a species group there is greater similarity, the divergence between phylogenetically more distant species is variably greater for different domains of the *hsr*ω gene.

### Search for hsrω homologs in other Dipterans

Since the 5’ and 3’ splice junctions of the *omega* intron are ultra-conserved in *Drosophila* while the *miR4951* is also present in nearly all *Drosophila* species, we searched for presence of these sequences in genomes of other dipterans whose sequences are available at NCBI. A 3236bp long scaffold (LWKS01024471.1) in *Zaprionus indianus* genome (Khanna & Mohanty, 2017) showed presence of the 60nt long *omega* intron’s 3’-splice junction sequence. However, this scaffold did not include any region with homology to the *omega* intron’s conserved 5’-splice junction sequence. The 16nt 5’ splice junction sequence did not find any significant match anywhere else in the publicly available genome sequence of *Zaprionus indianus*. Search for *miR4951* in the *Zaprionus indianus* genome sequence revealed high sequence homology in a 4239nt long scaffold (LWKS01014929.1). However, none of these two scaffolds showed sequence homology with other parts of the *hsr*ω gene. These two scaffolds did not show overlap with any other genomic sequence scaffolds available for this species. Therefore, it could not be ascertained if the 60 nt homologous sequence in the LWKS01024471.1 scaffold of *Zaprionus indianus* is part of *hsr*ω-like gene.

None of the other dipteran species genomes showed presence of sequences homologous to either the 5’ or 3’ splice junctions of the *omega* intron. However, earlier cytogenetic evidences for presence of tandem repeats and sequence divergence at one of the heat shock induced Balbiani rings in different species of *Chironomus* (Botella *et al*., 1991; Martinez-Guitarte *et al*., 2008; MartínezL□Guitarte & Morcillo, 2014) and the binding of Hsp83 at such a puff site following heat shock (Morcillo *et al*., 1993) suggested that this locus may be a functional homolog of the *Drosophila hsr*ω gene in *Chironomus*. Likewise, one of the heat shock puffs in the mediterranean fruit fly *Ceratitis capitata* has been reported (Semeshin *et al*., 1995) to harbour giant electron dense ribonucleoprotein (RNP) granules, similar to the omega speckles seen at the *hsr*ω locus and in nucleoplasm (Dangli *et al*., 1983; Prasanth *et al*., 2000). Thus it appears that a functional equivalent of the *hsr*ω locus may exist in other dipterans as well.

## Discussion

Early cytogenetic and molecular studies indicated that the non-coding *93D* or the *hsr*ω locus of *D. mel* is functionally conserved in diverse species of *Drosophila* with a similar genomic architecture but rapidly diversifying base sequence (Garbe *et al*., 1986; Garbe *et al*., 1989; Garbe & Pardue, 1986; Lakhotia, 1989; Lakhotia & Singh, 1982). Therefore, in our study, the base sequences of different regions were compared individually in the 34 species belonging to different lineages within the *Drosophila* genus. This has provided better insights into evolutionary trends of this large lncRNA gene.

Several of the species that were examined in earlier cytogenetic studies for presence of the benzamide and heat shock inducible orthologue of the *93D* locus of *D. mel* (Table 1) are also included in the present bioinformatic search for the *hsr*ω orthologue. In all cases, the present bioinformatic analysis identified the same loci that were identified earlier as the *93D* puff orthologue. This confirms that the genomic sequences retrieved by us are indeed orthologues of the *hsr*ω gene of *D. mel*. This is further substantiated by the finding that in 31 of the 34 species examined in this study, the genes flanking the retrieved putative *hsr*ω sequences were the same. The remarkable synteny of the *hsr*ω locus with its flanking genes is important since the 8 MB region surrounding this set of genes is associated with the *In(3R)Payne*, a common cosmopolitan inversion in *D. mel* whose polymorphism is correlated with geographic clines and, therefore, this region has also been referred to as a ‘*supergene*’ (Anderson *et al*., 2003; Durmaz *et al*., 2018; Kennington *et al*., 2006). It is significant that the genomic region associated with *In(3R)Payne* is also associated with adaptive clinal inversion polymorphism in *D. sub* and *D. buzzatii* populations (Hoffmann *et al*., 2004). Since the *hsr*ω is an important member of this ‘supergene’ complex in conferring thermal adaptation in clinal populations of *D. mel* (Collinge *et al*., 2008; McColl *et al*., 1996; McKechnie *et al*., 1998; Rako *et al*., 2007), the observed commonality of the flanking genes in other species of *Drosophila* may indicate comparable roles for the *hsr*ω orthologue in thermal and other adaptive features in other species as well. Our inability to identify these genes to flank the putatively identified *hsr*ω orthologue in *D. ame, D. bus* and *D. nas* is likely to be due to limitations of the genome sequence as presently available, although at this point some genomic rearrangements in these species during their diversification cannot be ruled out. Incomplete curation and/or inaccuracy in the assembly of genome sequences may also account for the identification of more than one *hsr*ω orthologue in some species and for our inability to assemble a contiguous stretch representing the *hsr*ω gene sequence in species like *D. eug, D. moj, D. nas, D. obs* and *D. rho*. Our analysis also suggests that the *GD15087* in *D. sim* is orthologue of *mod* and the annotation of *mod* orthologue.

Our study shows that regions flanking the *omega* intron’s 5’ and 3’ splice junctions are ultra-conserved in the 34 *Drosophila* species examined in this study. Our preliminary BLAST analysis (data not presented) suggests that the ultra-conserved exon-intron junctions of the *omega* intron are present in other *Drosophila* species whose genome sequences has now become available at NCBI. Intriguingly, unlike the generally observed greater evolutionary divergence of intronic than of exonic sequences, we found that the *omega* intron’s sequence divergence in different *Drosophila* species paralleled their phylogenetic relationship (Bai *et al*., 2007; O’Grady & DeSalle, 2018), while sequences of other regions (including the exons, other unique regions and tandem repeats associated with *hsr*ω gene) showed greater divergences, which do not follow the known phylogenetic relationship of these species. The unusual ultra-conservation of extended sequences flanking the *omega* intron splice junctions is unlikely to be related to general splicing functions since these appear unique to the *omega* intron with no other gene in any of the *Drosophila* species carrying a combination of the 16 bp 5’ and 60 bp 3’ splice junctions. Although 82 other genes in the examined *Drosophila* species carried 5’-splice junction sequences that were 69% to 87% similar to the 16 bp long ultra-conserved 5’ splice junction of the *omega* intron, none of these genes carried any similarity at their 3’-spice junctions with the conserved 60 bp at *omega* intron’s s’-splice junction. Since any obvious functional commonality between these 82 genes was not apparent, the significance of similarity with the *omega* intron’s 5’ junction sequence in these 82 genes is not clear. It is also intriguing that 50% of these occurrences showed similarity in antisense orientation. This aspect needs further analysis.

We searched for the potential of the conserved omega intron for encoding miRNA or snoRNA (data not presented) but did not find any such potentiality. Another possibility is that high conservation at intron junctions may be a feature of lncRNA sequences. However, our search for the splice junction sequences in several *Drosophila* lncRNAs *(roX1, roX2, yar, bxd, CR30121, CR30267, CR32835, CR44807, CR44807 and CR44810)* in the 34 different *Drosophila* species revealed that none the analysed lncRNA genes in different species showed comparable ultra-conservation of sequences flanking intron junctions (Supplementary Table S5). Therefore, we believe that the uniquely ultra-conserved *omega* splice junction flanks are essential for functionality of these ncRNAs by providing binding sites for some regulatory molecules and/or for generating some specific unique structural features. They may also help in the reported high stability of this intronic RNA (Garbe & Pardue, 1986).

It is interesting that the ultra-conserved *omega* intron’s 3’ splice junction, with full sequence identity, and *mir4951*, with significant similarity, were found in two genomic scaffolds of *Zaprionus indianus* (Khanna & Mohanty, 2017) a close relative of *Drosophila*. Since within the *Drosophila* genus we did not find this junction sequence or the *miR4951* to be present in any other gene with such high sequence identity, it is tempting to speculate that the genomic scaffold which harbors the *omega* intron’s 3’ splice junction sequence may be part of the *hsr*ω gene in *Zaprionus indianus*. However, in the absence of complete and assembled genome for *Zaprionus indianus*, this possibility cannot be confirmed at this stage.

A previous cytological search for the *93D* like locus (Nath & Lakhotia, 1991) suggested absence of a benzamide, colchicine, vitamin B6 or heat shocked glands homogenate inducible locus in *Anopheles* and *Chironomus* species. Our present search for the ultra-conserved *omega* intron splice junctions in other dipteran genomes also failed to identify a definitive *hsr*ω orthologue. Nevertheless, since as noted above one of the heat shock induced loci in different species of *Chironomus* and in the mediterranean fruit fly *Ceratitis capitata* appears functionally similar to the *hsr*ω locus in several respects like rapidly evolving tandem repeats, presence of large RNA particles, Hsp83 binding etc (Botella *et al*., 1991; Martinez-Guitarte *et al*., 2008; MartínezLGuitarte & Morcillo, 2014; Morcillo *et al*., 1993; Semeshin *et al*., 1995), it seems likely that a functional equivalent of the *hsr*ω gene of *Drosophila* is present in diverse dipterans as well. Multi-pronged and comprehensive cytogenetic, cell biological and genomic approaches are needed to confirm this possibility. Functional similarity between the cell stress-inducible human *Sat-III* and the *hsr*ω transcripts (Chung *et al*., 2018; Jolly & Lakhotia, 2006) further suggests that gene loci producing lncRNAs that are functionally analogous to *hsr*ω transcripts indeed exist in diverse taxa.

Our promoter analysis suggested that the promoter at TSS2 of *D. mel* is the major functional promoter as this shows higher conservation and most of the *hsr*ω transcripts in this and the other species, whose transcriptomic information is available at the FlyBase, are initiated from this site. Our bioinformatic analysis predicted existence of the TSS1 promoter in corresponding region in most other species, although the current annotations of *hsr*ω transcripts in *Drosophila* species do not show presence of a functional TSS1 except in *D. mel*. It may be noted that the TSS1 in *D. mel hsr*ω gene was identified at the FlyBase only recently following in-depth RNA-seq studies. In view of this, we believe that with more detailed transcriptomic studies, existence of TSS1 is likely to be uncovered in other species as well. Since the TSS2 and *mir4951* sequences are present at proximal and distal ends, respectively, of the *hsr*ω orthologue in nearly all the *Drosophila* species examined and since TSS1 is also likely to be present at the most proximal end of *hsr*ω gene in different species, we suggest that the *hsr*ω orthologue in different species of *Drosophila* start with TSS1 and terminate after *mir4951* as in *D. mel* (Figs. 4 and 5).

The present *in silico* analysis failed to definitively predict the transcription termination sites, which must exist in multiple places in view of the known multiple transcripts produced by this gene in different species. This failure may be related to the possibility that these transcripts may not be polyadenylated and, consequently, they may not carry the typical signatures required by the different online tools used for TTS prediction. More exhaustive transcriptomic data are needed to identify the expected multiple TTS sites.

The tandem repeats uniquely associated with the *hsr*ω gene in *D. mel* have attracted considerable attention due to their specific association with the omega speckles (Lakhotia, 2011; Prasanth *et al*., 2000; Singh & Lakhotia, 2015). In agreement with results of the initial cloning and sequencing of the *hsr*ω gene from three species in the 1980s (Garbe *et al*., 1986; Peters *et al*., 1982; Peters *et al*., 1984; Ryseck *et al*., 1987), the present bioinformatic analysis also revealed that the *hsr*ω gene associated repeats are the most variable component in spite of their being unique to this gene. The rapid divergence between the tandem repeats associated with *hsr*ω gene in different species is in general agreement with the widely known rapid diversification and concerted evolution of repeat sequences during speciation (Chen *et al*., 2016; Chodroff *et al*., 2010; Corona-Gomez *et al*., 2019; Haerty & Ponting, 2013; Kapusta & Feschotte, 2014; Lakhotia, 2017a; Lakhotia, 2017b; Ponting, 2017; Tavares *et al*., 2019; Zhang *et al*., 2017), although unlike many other genomic repeats, the *hsr*ω repeats have not spread in the genome.

The rapid diversification of the *hsr*ω associated tandem repeats within a species, and more so between species, is intriguing in the context of their essential role in organization of omega speckles in *D. mel* which dynamically regulate the functional availability of various hnRNPs and related proteins (Lakhotia, 2011; Prasanth *et al*., 2000; Singh & Lakhotia, 2015). Since as in *D. mel*, the various hnRNPs also accumulate at the *hsr*ω gene site in stressed cells in other species that have been examined so far (Chowdhuri & Lakhotia, 1986; Dangli *et al*., 1983), we believe that the nuclear transcripts of the *hsr*ω orthologues perform similar functions in other *Drosophila* species as well. Apparently, the tandem repeats of the *hsr*ω gene discharge their common functions despite the rapid sequence divergence. Interestingly, the Human repetitive *Sat III* sequences also do not show any similarity with tandem repeats of the *hsr*ω gene but perform comparable functions (Chung *et al*., 2018; Jolly & Lakhotia, 2006). As known for many lncRNAs, especially for those associated with membraneless phase-separated nuclear sub-structures, the secondary and higher order structures attained by the specific lncRNA orthologue are critical (Lakhotia, 2017a; Lakhotia, 2017b; Lin *et al*., 2018). The earlier (Garbe *et al*., 1986; Garbe *et al*., 1989; Peters *et al*., 1984; Ryseck *et al*., 1987) and our current finding of high conservation of a nonamer sequence within the poorly conserved *hsr*ω gene associated tandem repeats in different species is significant in this context. The heterogeneous RNA binding proteins (hnRNPs) like Hrb87F, Hrb98DE, which are integral parts of the omega speckles (Lakhotia, 2012; Prasanth *et al*., 2000; Singh & Lakhotia, 2015), have binding affinity for the *hsr*ω repeat associated nonamer and related sequences (Zu *et al*., 1998). Thus despite the rapid divergence of tandem repeats, presence of multiple copies of classical or variant nonamers in the middle region of the larger *hsr*ω transcripts may provide a basis for regulation of hnRNP dynamics by the *hsr*ω transcripts in different species. Present analysis suggests that while the basic repeat unit could get rapidly altered, there has been a strong purifying selection for retaining the nonamer sequences. It would be interesting to examine if these nonamers indeed provide ‘landing sites’ for one or more of the hnRNPs during biogenesis of omega speckles which occurs as these transcripts are being produced at the *hsr*ω gene locus (Singh & Lakhotia, 2015). Examination of status of omega speckles in different species and their dynamics under different conditions of cell stress, including in those that show fewer or no tandem repeats as part of the *hsr*ω gene, would be very interesting.

Earlier analysis of the ORFω in three distantly related species (*D. mel, D. hyd* and *D. pse*), suggested this to be rather poorly conserved (Fini *et al*., 1989; Garbe *et al*., 1986). Our present analysis using data from closely and distantly related species, however, revealed a relatively higher conservation in context of sequence and secondary structure of the ORFω in the 34 examined species. The first four residues at the amino terminal sequence in 27 species were MEKC/MKKC/MQKC/MQMC/MEMC while in others the additional residues in this region were seen in an evolutionary conserved pattern. It is interesting that in species belonging to the *obscura* group, the extra residues at the amino-terminal were MY/HIYCSTAM while in the *ananassae* group (*D. ana* and *D. bip*), a Y was seen after the first methionine residue. These may reflect lineage-specific insertions. The carboxyl terminal of this peptide showed two principle motifs, viz., “G/TPT” in 21 species belonging to the closely related *melanogaster* and *obscura* groups and “RRLK” in 12 species belonging to *Dorsilopha, immigrans, Hawaiian* and *virilis* groups. *D. wil*, which is a separate clade (O’Grady & DeSalle, 2018) carries very different residues (AVA) at the carboxy terminal of the Omega peptide. Thus the different motifs at the carboxy terminal of this small peptide follow the phylogenetic diversification in the *Drosophila* genus during the past 60-70 million years (Bai *et al*., 2007; O’Grady & DeSalle, 2018). Such conservation of amino and carboxy-terminal sequences and its generally conserved secondary structure indicates potential functionality of the Omega peptide. The present *in silico* structural analysis suggests that the Omega peptide can form an alpha helical structure, which agrees with recent evidences for the existence of alpha helical short peptides (Baeriswyl *et al*., 2019; Bystroff & Garde, 2003; Kelso *et al*., 2004; Shin & Hahm, 2004). The first four N-terminal residues, known as capping box sequences, are considered to be important for the stabilization of alpha-helices in proteins and peptides (Aurora & Rosee, 1998; Forood *et al*., 1993; Harper & Rose, 1993; Krstenansky *et al*., 1989; Marqusee & Baldwin, 1987). It appears that the N-terminal conserved MEKC/MKKC/MQKC/MQMC/MEMC sequences may function as a potential N-Cap for the short omega peptide. The carboxy terminal motifs (Schellman motif and αL motif) often include Glycine residues at the C- cap sequences (Aurora *et al*., 1994). The GPT motif, seen in the Omega peptide in many species, could, therefore, be a potential C-box for the predicted alpha helix. Significance of the RRLK sequence at the Omega peptide C-terminus in other species remains unclear. Its functionality is supported by translatability of the *D. mel* ORFω *in vitro* (Fini et al., 1989) as well as *in vivo* (R. K. Sahu and S. C. Lakhotia, unpublished). Further support for functional requirement of the Omega peptide is provided by our other finding that CRISPR/Cas mediated targeted deletion of the ORFω results in several defects, including complete sterility of females (R. K. Sahu, Rima Saha and Lakhotia, unpublished). These aspects are being examined further.

The evolutionary relationships of the *hsr*ω gene revealed by its sequence comparison in different species generally agree with the previously published phylogenies of different *Drosophila* species (Bai *et al*., 2007; O’Grady & DeSalle, 2018; Seetharam & Stuart, 2012; Seetharam & Stuart, 2013; Van Der Linde *et al*., 2010). However, as noted above, different domains of this gene have diverged at varying rates even within a single species group/sub-group. Present analysis of the *hsr*ω orthologues in 34 species of *Drosophila* agrees with an earlier finding based on comparison of this gene’s base sequence in three species (Garbe *et al*., 1989) that while the proximal part, encompassing exon1, the *omega* intron and exon 2 of the *hsr*ω gene is better conserved in different species, the distal part is more variable. The higher variability of the distal part is mostly due to the rapidly diversifying tandem repeats, which seems to be further compounded by the incomplete genomic information for this region in different species. It is expected that as more genomic and transcriptomic data become available, the architecture of the *hsr*ω lncRNA gene in different species would be better known. Notwithstanding these limitations at this time, the similar organization, amidst sequence variability, of the two exons, *omega* intron and the ORFω is remarkable and indicates need for greater attention to this region for its functional significance.

It is generally recognized that compared to the relatively high sequence conservation in protein-coding and a few non-coding genes due to purifying selection, majority of the lncRNA genes show little conservation in their transcribed sequences except for small domains (Chen *et al*., 2016; Lakhotia, 2017a; Lakhotia, 2017b) (Chodroff *et al*., 2010; Corona-Gomez *et al*., 2019; Haerty & Ponting, 2013; Kapusta & Feschotte, 2014; Ponting, 2017; Tavares *et al*., 2019; Zhang *et al*., 2017). Our analysis of the nucleotide sequence of the relatively large *hsr*ω lncRNA gene in the genus *Drosophila* revealed a peppered sequence conservation amidst a similar architecture. Like the nuclear transcripts of the *hsr*ω gene which organize the omega speckles (Prasanth *et al*., 2000; Singh & Lakhotia, 2015), the large NEAT1 lncRNAs organize nuclear paraspeckles in different mammals, despite poor sequence conservation. It is suggested that the NEAT1 transcripts attain a unique structure because of long-range RNA interactions and thus provide appropriate scaffold for the assembly of paraspeckles (Lin *et al*., 2018). It would be interesting to examine structural features of the various *hsr*ω transcripts in different species to see if, in spite of the general sequence variability, conserved domains like the 5’ and 3’ *omega* intron flanking sequences and/or the conserved nonamers help in attaining certain higher order structure/s that provide the required framework for assembly of the omega speckles. Likewise, further studies are necessary to understand the functional significance of the short ORFω, which although present in all species, also shows limited sequence conservation,

Our present annotation of the unusual *hsr*ω gene, whose multiple transcripts perform diverse functions as lncRNAs, architectural RNAs and through production of a small peptide (Lakhotia, 1987; Lakhotia, 2011; Lakhotia, 2017a), would be very useful for further studies on this gene’s evolution and functions in different species of *Drosophila*. The extensive sequence diversification amidst a rather conserved architecture of the *hsr*ω lncRNA gene, which is essential for organism’s survival under normal as well as stress conditions, makes this locus a uniquely interesting model for studying evolutionary changes and factors that shape such changes.

## Acknowledgements

We thank the Department of Biotechnology, Ministry of Science and Technology, Govt. of India, New Delhi for supporting this work (BT/PR6150/COE/34/20/2013). SCL is currently supported by the Science & Engineering Research Board (SERB), Govt. of India, as SERB Distinguished Fellow. RKS is supported by the Council of Scientific and Industrial Research, New Delhi through research fellowship. EM was supported by Summer Project fellowship of the Science Academies.

## Author contributions

RKS, EM and SCL undertook the bioinformatic analyses and wrote the manuscript.

